# Diffusion of activated ATM explains γH2AX and MDC1 spread beyond the DNA damage site

**DOI:** 10.1101/2023.10.02.560532

**Authors:** Georgi Danovski, Greta Panova, Bradley Keister, Georgi Georgiev, Krastan B. Blagoev, Stoyno S. Stoynov

**Affiliations:** Institute of Molecular Biology, Bulgarian Academy of Sciences, 21, G. Bontchev Str., Sofia 1113, Bulgaria; Department of Mathematics, USC, CA 90089, USA; Department of Physics, UCSD, CA 92093, USA; Faculty of Mathematics and Informatics, Sofia University, St. Kliment Ohridski, 5 James Bourchier Blvd., 1164 Sofia, Bulgaria; National Science Foundation, Alexandria, VA 22230, USA; Department of Biophysics, Johns Hopkins University, Baltimore, MD 21218, USA; Institut Curie, PSL Research University, Sorbonne Université, CNRS UMR3664, Paris, France

**Keywords:** DNA damage response, ATM, MDC1, cohesion, NIPBL, RAD21, γH2AX

## Abstract

During DNA repair, ATM-induced H2AX histone phosphorylation and MDC1 recruitment spread megabases beyond the damage site. The mechanism underlying this spread remains unclear. To elucidate this step of the DNA damage response, we measured ATM and MDC1 recruitment kinetics at micro-irradiation-induced complex DNA lesions. While MDC1 spreads beyond the damage-induced focus of ATM accumulation, cohesin loader NIPBL and cohesin subunit RAD21 accumulate later than MDC1. Such delayed recruitment suggests that, in complex DNA lesions, loop extrusion cannot alone explain γH2AX and MDC1 spread. To determine if diffusion of activated ATM could account for the observed behavior, we measured the exchange rate and diffusion constants of ATM and MDC1 within damaged and unperturbed chromatin. Based on these measurements, we introduced a quantitative model that describes the dynamic behavior of ATM and subsequent MDC1 spread at complex DNA lesions. Altogether, we propose activated ATM diffusion as an underlying mechanism of MDC1 spread.

## Introduction

Cell function depends on a myriad of interdependent processes organized in complex biochemical networks, involving chemical interactions and the transport of biomolecules inside the cell. As a result, various macromolecules are heterogeneously distributed among cellular compartments, and their concentrations vary in space and time during the cell cycle. The DNA damage response is one such complex biochemical network, involving more than 300 proteins that participate in complex biophysical interactions to form interconnected pathways. In addition to the well-established consequences of coding region mutations on protein function, alterations in regulatory regions influence repair factor concentration and may compromise repair proficiency, leading to genomic instability and cancer.

A comprehensive understanding of the DNA damage response would require not only knowledge of the proteins involved and their interactions, but also of their spatiotemporal dynamics within the cell. To this end, one needs precise measurements and modeling of repair factor kinetics. We and others have measured the kinetics of recruitment and release of repair factors at DNA damage sites, an approach that allows us to obtain a detailed quantitative understanding of DNA repair mechanisms through mathematical modeling of the physical processes taking place ^1,2^.

A key event during the repair of DNA double-strand breaks (DSB) is the phosphorylation of the histone H2AX on serine 139 (γH2AX)^3-6^ by the ataxia telangiectasia modified (ATM) protein kinase. This modification is subsequently recognized by the mediator of DNA damage checkpoint protein 1 (MDC1) ^7-10^. While ATM and the MRN complex, which recruits the former, both localize to the DNA damage site, γH2AX spreads out over several megabases of DNA beyond the damage site.

Recently, using structure illuminated super-resolution microscopy in fixed cells, Natale et al. (2017) showed that individual large repair protein foci consist of several nano-foci organized in clusters around a DSB. These clusters were suggested to depend on chromatin architecture and CTCF. Using 4C-seq ^11^, found that the spatial distribution of γH2AX is correlated with the chromatin contacts near DSBs. Based on circular chromatin conformation capture coupled to 4C-seq, Legube et al.^12^ also proposed loop extrusion as a major determinant of γH2AX distribution. That is, cohesin-dependent loop extrusion at the DSB drives ATM-mediated H2AX phosphorylation at the megabase scale. Despite extensive studies into the matter, the exact mechanism through which the H2AX phosphorylation and downstream MDC1 recruitment spread beyond the repair site remains unclear.

In the present study, we demonstrate that ATM is not anchored to chromatin at micro-irradiation (micro-IR)-induced complex DNA lesions, but frequently binds and unbinds from the damage site. Furthermore, we found that MDC1 recruitment kinetics is faster than those of cohesin loader NIPBL and cohesin subunit RAD21. These findings suggest another mechanism of H2AX phosphorylation beyond the damage site. Our precise measurements of ATM and MDC1 recruitment and exchange kinetics, together with the data on their spatiotemporal concentrations, allow us to test several reaction-diffusion mathematical models^13^ for the kinetics of γH2AX and MDC1 spread. The experimental measurements coupled to mathematical modeling demonstrate that activated ATM diffusing away from the DNA damage site can explain the observed spatial and temporal distribution of γH2AX and MDC1.

## Results

### Dynamics of MDC1 at complex DNA lesions

To follow H2AX phosphorylation in space and time in living cells, we measured the kinetics of MDC1 at a complex DNA lesion, generated through micro-IR within a small, localized three-dimensional region. To this end, we used an EGFP-tagged MDC1 transgenic HeLa Kyoto cell line^14^, generated through bacterial artificial chromosome (BAC) recombination^15^. In this line, the tagged MDC1 is expressed at near-physiological levels ^16^ under cell-cycle control. Our results (Figures 1A and 1B; Movie S1) show that MDC1 is quickly recruited to the site of micro-IR^1,17^, with a half-time of 55s, reaching its maximum levels at around 900s, whereafter it spreads around the complex lesion. As a result, MDC1 is heavily depleted from other regions of the nucleus following damage induction. Comparing the kinetics of MDC1 recruitment and depletion, we see a significant delay in the latter process, most likely caused by the complex transport of MDC1 along chromatin, which differs from simple diffusion.

**Figure 1.**
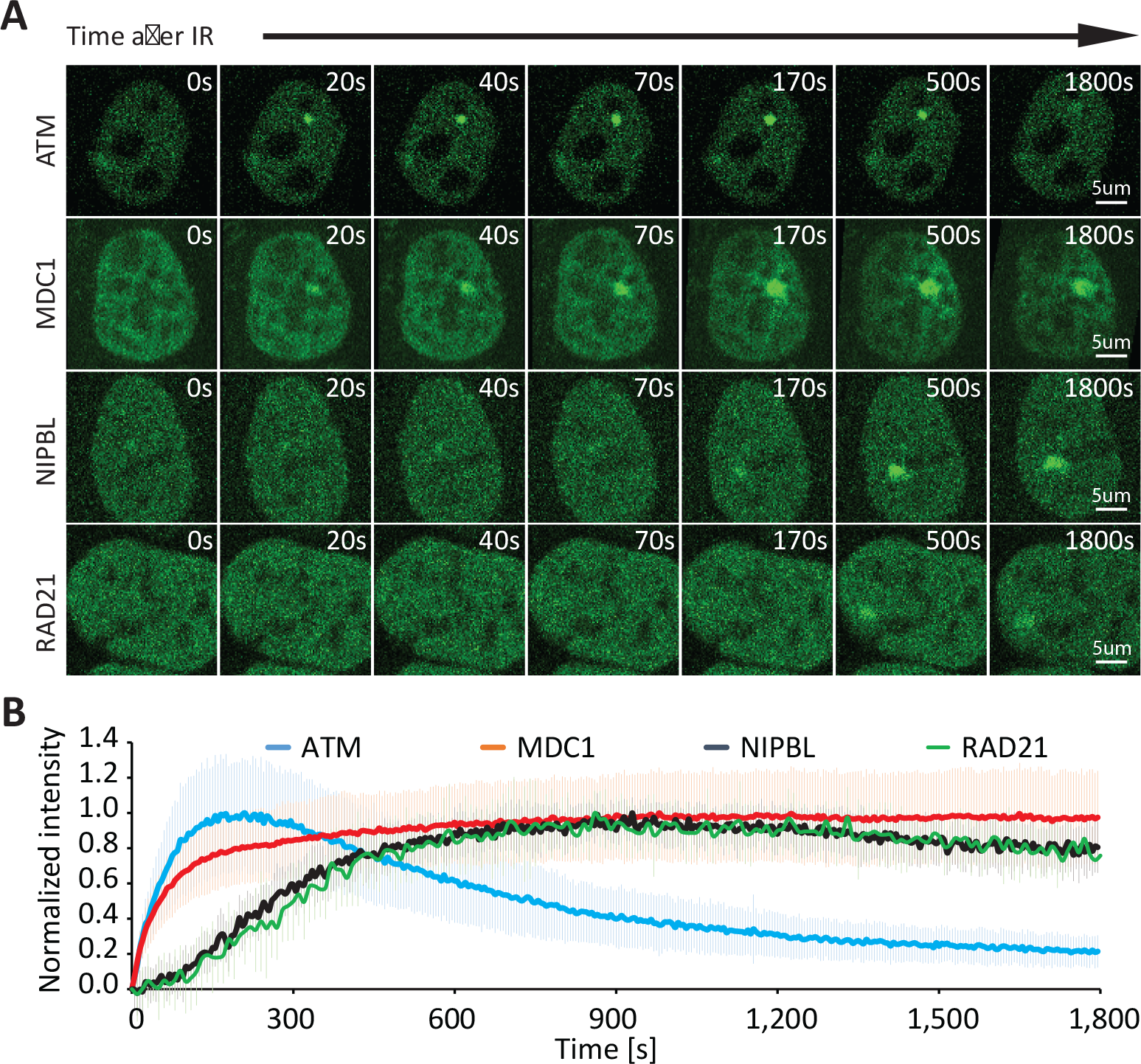
Spatiotemporal dynamics of ATM, MDC1, mNIPBL, and RAD21 at sites of complex DNA damage. All time points can be observed in Movies S1 and S3. (A) Representative time-lapse microscopy images of the spatial distribution of ATM, MDC1, mNIPBL, and RAD21 after the induction of a complex DNA lesion. (B) ATM, MDC1, mNIPBL, and RAD21 recruitment kinetics at the sites of complex DNA damage (error bars show the standard deviation).

Fluorescence recovery after photobleaching (FRAP) analysis showed that the kinetics of MDC1 exchange at the damage site was also slow (with a half-time of 27.5±11s) (Figure S1A and S1B; Movie S2). We performed further FRAP experiments to study MDC1 mobility in the absence of damage, observing MDC1-EGFP signal recovery after bleaching a small region of the nucleus (Figure S1C). MDC1 mobility was approximately an order of magnitude slower (average D=0.055±0.033μm^2^/s) than expected for the pure diffusion of a 200kDa (0.4-0.6μm^2^/s) protein. It is known that MDC1 is weakly bound to the chromatin even in the absence of DNA damage ^17,18^. This can also be inferred from the irregular spatial distribution of MDC1, which resembles the spatial structure of chromatin. Such an association of MDC1 with chromatin, in the absence of a lesion, could explain its slower effective diffusion.

### Chromatin binding affects MDC1 mobilization to a secondary damage site

We reasoned that the slow mobility of MDC1 may have an impact on DNA repair at a second DNA damage site. To interrogate this, we induced a second lesion at a distant nuclear site 30 min after the initial micro-IR insult. Second micro-irradiation led to decrease of the amount of MDC1 at the first damage site. (Figure 2A and 2B). Further, kinetic of the removal of MDC1 at the first focus is significantly slower than the kinetic of the recruitment of the MDC1 at the second lesion. These results demonstrate that the limited MDC1 concentration and its slow transport in the nucleus due to chromatin binding influence the kinetics and spread of MDC1 in the presence of two consecutive micro-IR-induced damage foci.

**Figure 2.**
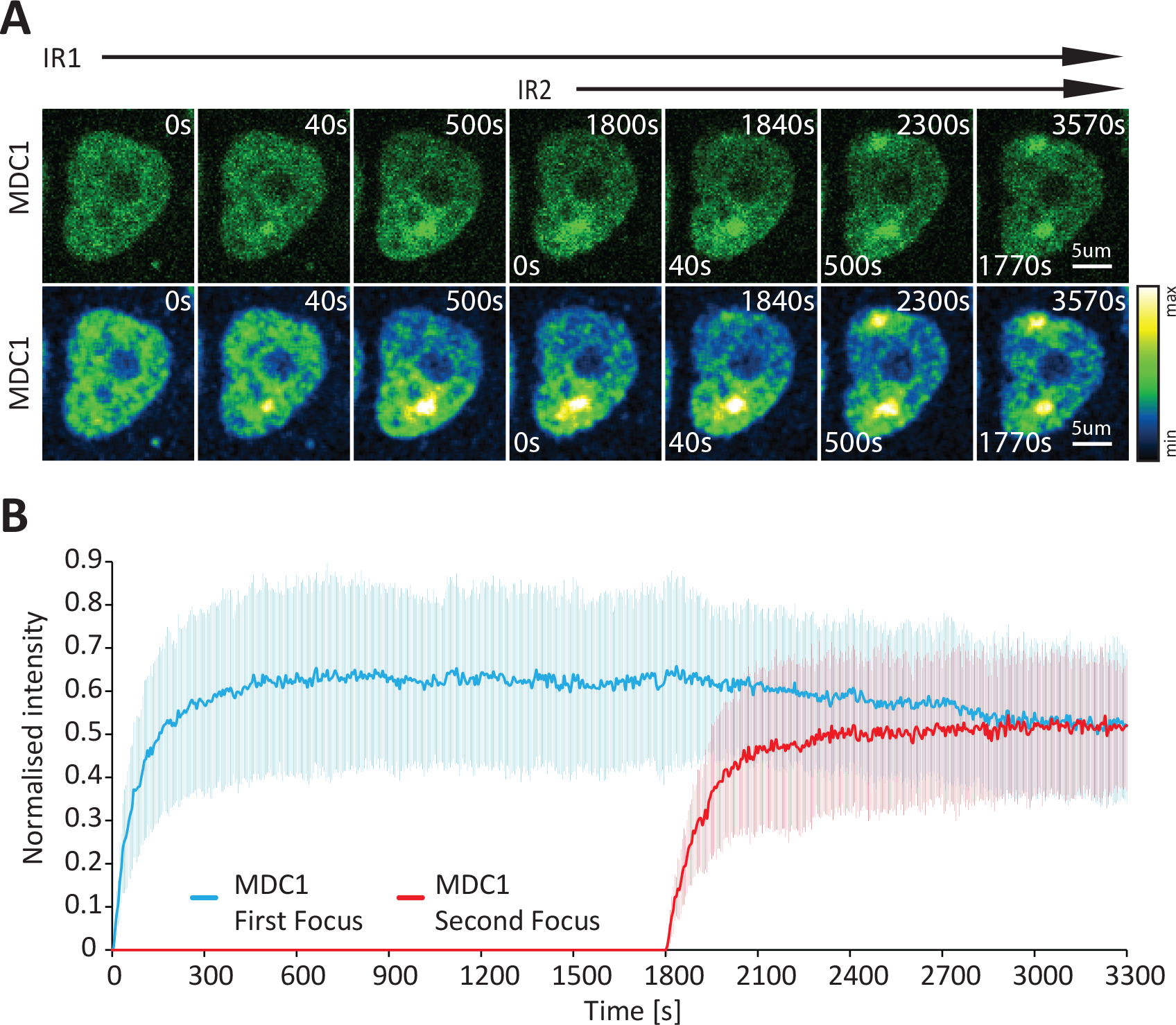
MDC1 recruitment kinetics at two consecutive micro-irradiation-induced complex DNA lesions. (A) Representative time-lapse images are presented in the row above. The row below presents the same images where intensity is color coded. (B) Recruitment kinetics of consecutive micro-irradiation-induced MDC1 foci. (error bars show the standard deviation)

### ATM is rapidly exchanged at complex DNA lesions

To better understand MDC1 spread kinetics, we also measured the kinetics of ATM, the major apical kinase responsible for phosphorylating H2AX histones at DNA damage sites. ATM is recruited earlier (half-time of recruitment: 40s) than MDC1 ^1^ and, in contrast to the latter, accumulates only within the DNA damage site (Figure 1A and 1B; Movie S1). ATM rapidly accumulates upon micro-IR (reaching a maximum after 210s), which is followed by a rapid decrease shortly thereafter Figure 1B). The transition between ATM accumulation confined within the damaged region and the subsequent expansion, or spread, of γH2AX and MDC1 remains mechanistically undetermined.

Arnould and coworker put forth a model wherein activated ATM is localized to the boundaries of cohesin-dependent chromatin loops next to a DSB ^12^. Subsequently, accumulated repair machinery prevents loop extrusion in one direction, while extrusion proceeds in the other, with ATM phosphorylating the chromatin pulled through to spread γH2AX. ATM binding and phosphorylation while DNA is being pulled through suggests a rather slow ATM exchange. However, our FRAP measurements at micro-IR-induced complex lesions revealed very fast ATM exchange rates ^19^ (Figure S1A and S1B), close to those of freely diffusing ATM (Figure S1C)

### Late recruitment of loop extrusion machinery cannot account for MDC1 spread

To understand the time scale at which loop extrusion operates and thus validate the above-described model, we compared the accumulation rate of cohesin loader NIPBL, required for DSB-dependent cohesin recruitment and loop extrusion, with those of ATM and MDC1 (Figure 1A and 1B; Movie S1). This comparison showed that MDC1 and ATM recruitment kinetics are considerably faster than for mouse NIPBL (mNIPBL), which suggests that the MDC1 spread occurs before DSB-induced loop extrusion. This is also supported by the finding that MDC1 and RNF168 are required for mNIPBL loading ^20^. To directly confirm cohesin recruitment timing, we employed cells expressing endogenously tagged with mClover, cohesin subunit RAD21, which exhibits loop extrusion activity *in vivo* similar to the wild-type cells ^21,22^. RAD21 recruitment kinetics closely followed those of mNIPBL, being significantly slower than MDC1 (Figure 1A and 1B; Movie S3). These results strongly suggest that mechanisms other than loop extrusion may be responsible for γH2AX/MDC1 spread at complex DNA lesions.

An alternative explanation for γH2AX/MDC1 spread away from the micro-IR-induced DNA lesions is a model wherein the diffusing activated ATM phosphorylates chromatin along its path. In this model, ATM binds at the damage site, is activated, and then released. Upon its detachment from the damage site, freely diffusing activated ATM phosphorylates the chromatin along its path until inactivated. We introduced multiple quantitative mathematical models to describe this phosphorylation mechanism and confirm whether they are consistent with our measurements.

### Modelling of MDC1 spread based on diffusing activated ATM

A spatiotemporal mathematical model of physical processes should be able to quantitatively explain the kinetics of ATM and MDC1 recruitment and removal throughout the whole nucleus after induction of complex DNA lesions. The model should also faithfully describe the actual heterogeneous chemical reaction parameters derived from FRAP of ATM and MDC1 at DNA damage sites. The first model that we developed describes ATM and MDC1 data via the following reactions:

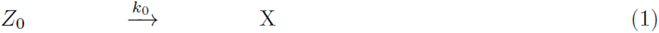

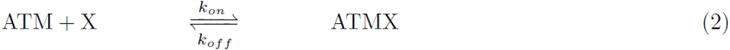

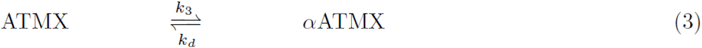

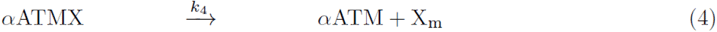

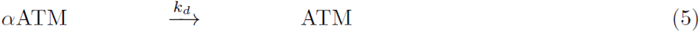

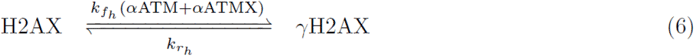

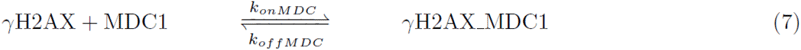

Here, Z_0_ represents the DNA damage, and X represents the proteins bound to damage sites, e.g. the MRN complex, which accumulates prior to ATM recruitment. ATMX represents the bound ATM, αATM is the activated ATM, αATMX represents the bound αATM and mX represents modified X after αATM removal. The partial differential equations describing these kinetic reactions for the corresponding concentrations are provided in the supplementary information (Data S1). We will call this model the “Standard αATM diffusion model”, or the “Standard model”, for simplicity (Figure 3). This model describes the DNA damage within the irradiated region of the nucleus, which are converted into a DNA damage-protein complex before the ATM binding, with a rate k_1_. ATM binds and unbinds to this complex with rates k_on_ and k_off_, respectively. ATM undergoes reversible activation at the damage site, whereafter activated ATM leaves the site irreversibly, with a rate k4. Consequently, the now freely diffusing activated ATM is deactivated with the same rate constant as the bound activated ATM. Once activated, both the bound and unbound αATM phosphorylate H2AX to γH2AX with a rate k_f_, which depends on the αATM concentration. Phosphorylated H2AX is then recognized by MDC1, forming a γH2AX/MDC1 complex.

**Figure 3.**
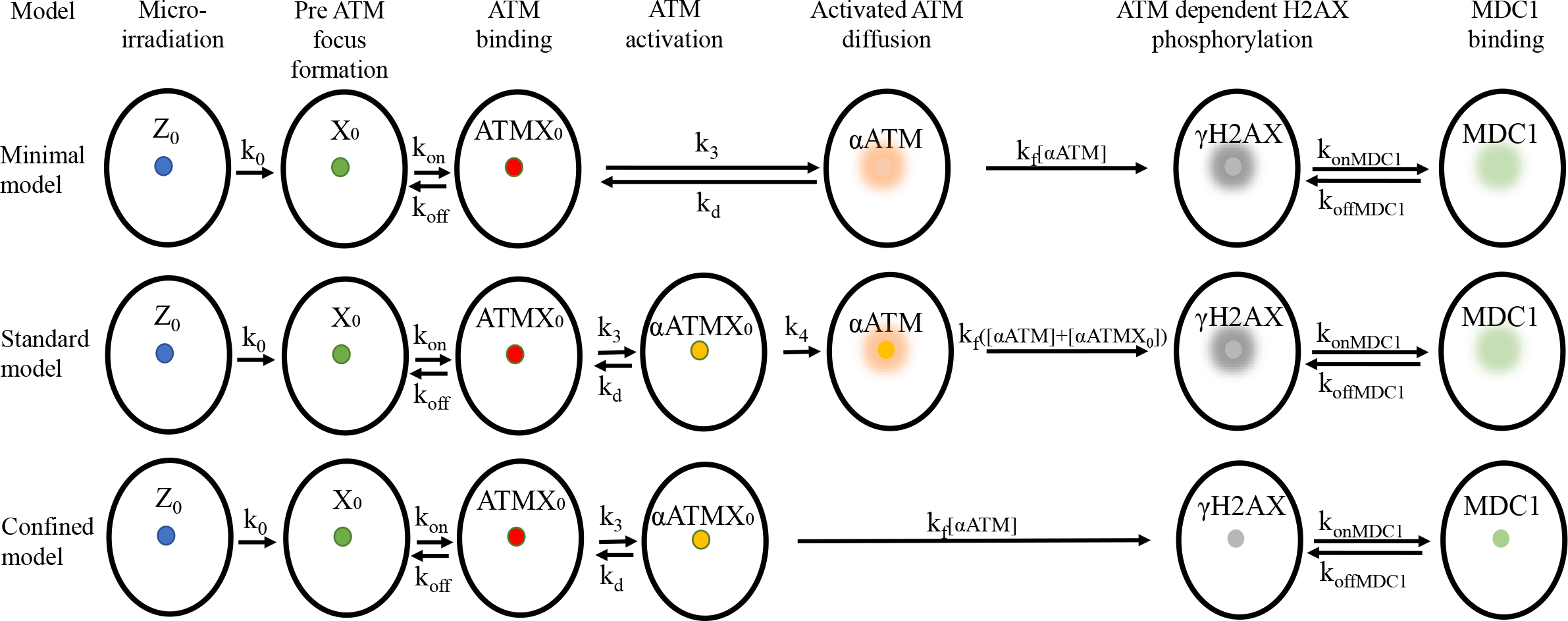
Graphical representation of the three mathematical models describing the ATM-dependent H2AX phosphorylation and MDC1 recruitment dynamics.

We model the biological processes shown in reactions (1-7) by including the diffusion of molecules through a system of reaction-diffusion partial differential equations for the corresponding concentrations. The reaction-diffusion equations that correspond to these reactions and the details of our numerical implementation are shown in the Supplementary information (Data S1). We created a dedicated software tool in order to simulate ATM and MDC1 nuclear dynamics following micro-IR as per an assigned model. The software enabled us to generate time-lapse images of the modelled process, in addition to two graphical representations. The first follows protein-of-interest concentration in a given nuclear region over time. The latter representation visualizes the spatial distribution of protein concentration within a given region as a time-lapse video. This tool enabled to compare experimental data against model predictions.

Using live-cell microscopy, we were able to measure several physical parameters that were implemented in the mathematical model (Table S1). The ATM diffusion constant was measured using FRAP of the unbound ATM (average D=0.664±0.34μm^2^/s). Complex MDC1 transport was modeled with an effective diffusion constant, which we also measured using FRAP of cells without DNA damage (average D=0.055±0.35μm^2^/s). This effective diffusion constant captures the free diffusion as well as the process of MDC1 binding and unbinding to the undamaged chromatin. Other input parameters were the ATM and MDC1 concentrations were calculated (material and methods) using their cell copy numbers, obtained in ^16^. We measured the time course of ATM and MDC1 fluorescence intensity at the DNA damage site as well as in a small region of interest outside this site. MDC1 spatial distribution around the damage site was also obtained. Using the initial ATM and MDC1 concentrations in the nucleus allowed us to convert the fluorescence intensity to concentrations.

To obtain the kinetic rates of reactions (1-9), we fit the standard model to our experimental data, which include: (1) the concentrations of MDC1 (Figure 4A and 4C) and ATM (Figure 5A and 5C) at the damage site as well as in a nuclear region away from it; (2) the concentrations of MDC1 (Figure 6A and 6B) and ATM (Figure 7A-C) after FRAP of the damage site. Importantly, in order to assess whether the proposed models can explain the observed pattern of MDC1 spread, we compare the measured spatial distribution of MDC1 (Figure 4B; Figure S2; Movie S4) and ATM (Figure 5B; Figure S3; Movie S5) around micro-IR-induced damage foci over time versus those predicted by the model.

**Figure 4.**
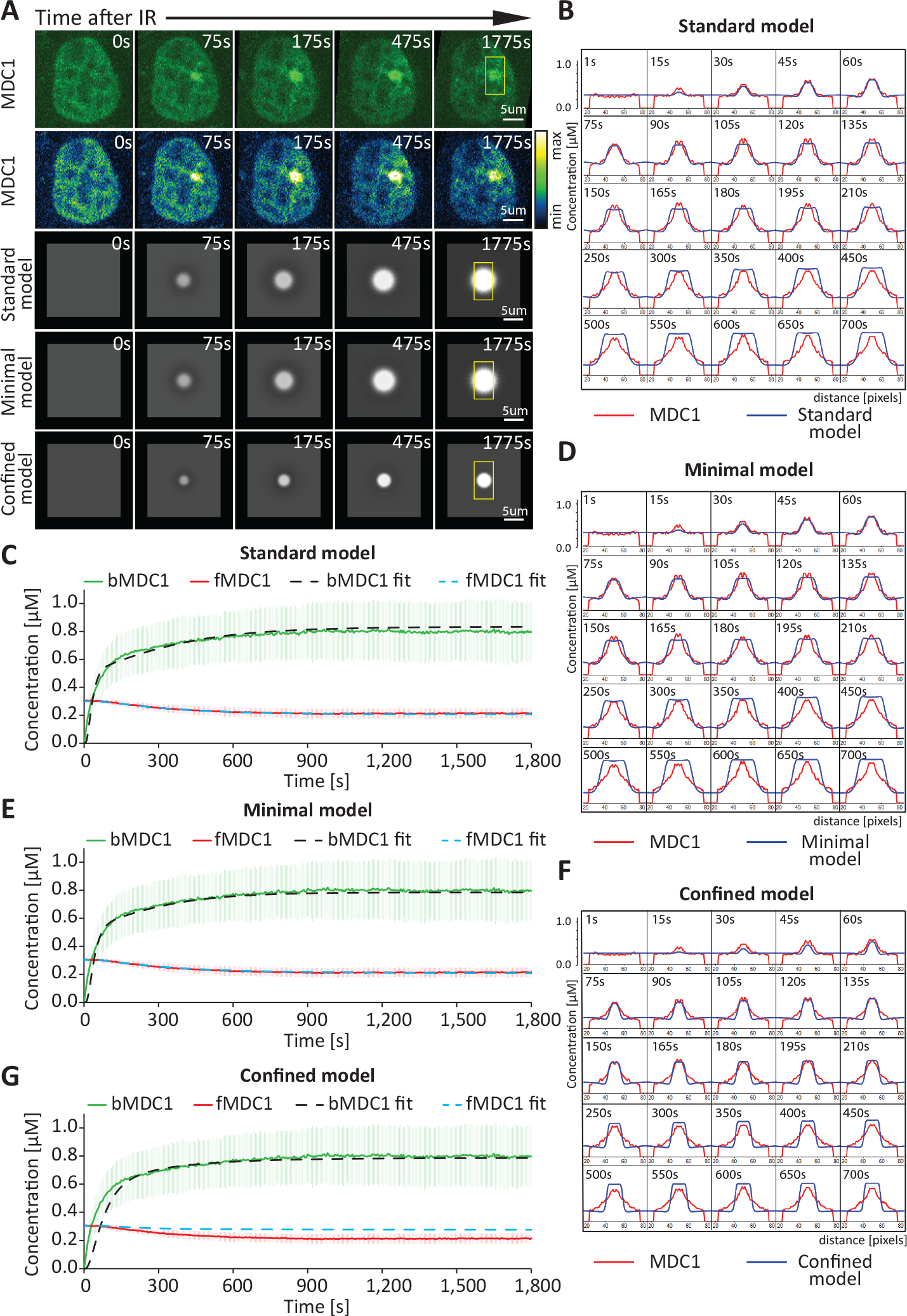
Comparison between the experimentally measured and predicted MDC1 dynamics at sites of complex DNA damage. (A) Comparison between time-lapse microscopy images and images generated via simulation based on the models of MDC1 distribution at sites of complex DNA damage. The yellow box covers a region of interest where the MDC1 spread presented in B, D, and F is measured. (B) Time-lapse representation of the MDC1 concentration profile across the micro-irradiation site (the region of interest in A). Red color indicates the measured MDC1 concentration spread. Blue indicates the best-fitting predicted concentration based on the standard model. (C) Graphical comparison between the experimentally measured MDC1 dynamics at sites of complex DNA damage and those predicted based on the standard model. Error bars represent the standard deviation. bMDC1 (bound MDC1), measured concentration (μM) of MDC1 within the DNA damage site; fMDC1 (free MDC1), measured concentration (μM) of MDC1 outside of the damage focus; bMDC1 fit, the best-fitting predicted concentration (μM) of MDC1 at the damage site, based on the standard model; fMDC1 fit, the best-fitting predicted concentration (μM) of MDC1 outside of the damage site, based on the standard model. (D) Time-lapse representation of the MDC1 concentration profile across the micro-irradiation site (the region of interest in A). Red color indicates the measured MDC1 concentration spread. Blue indicates the best-fitting predicted concentration based on the minimal model. (E) Graphical comparison between the experimentally measured MDC1 dynamics at sites of complex DNA damage and those predicted based on the minimal model. Error bars represent the standard deviation. bMDC1, measured concentration (μM) of MDC1 within the DNA damage site; fMDC1, measured concentration (μM) of MDC1 outside of the damage focus; bMDC1 fit, the best-fitting predicted concentration (μM) of MDC1 at the damage site, based on the minimal model; fMDC1 fit, the best-fitting predicted concentration (μM) of MDC1 outside of the damage site, based on the minimal model. (F) Time-lapse representation of the MDC1 concentration profile across the micro-irradiation site (the region of interest in A). Red color indicates the measured MDC1 concentration spread. Blue indicates the best-fitting predicted concentration based on the confined model. (G) Graphical comparison between the experimentally measured MDC1 dynamics at sites of complex DNA damage and those predicted based on the confined model. Error bars represent the standard deviation. bMDC1, measured concentration (μM) of MDC1 within the DNA damage site; fMDC1, measured concentration (μM) of MDC1 outside of the damage focus; bMDC1 fit, the best-fitting predicted concentration (μM) of MDC1 at the damage site, based on the confined model; fMDC1 fit, the best-fitting predicted concentration (μM) of MDC1 outside of the damage site, based on the confined model.

**Figure 5.**
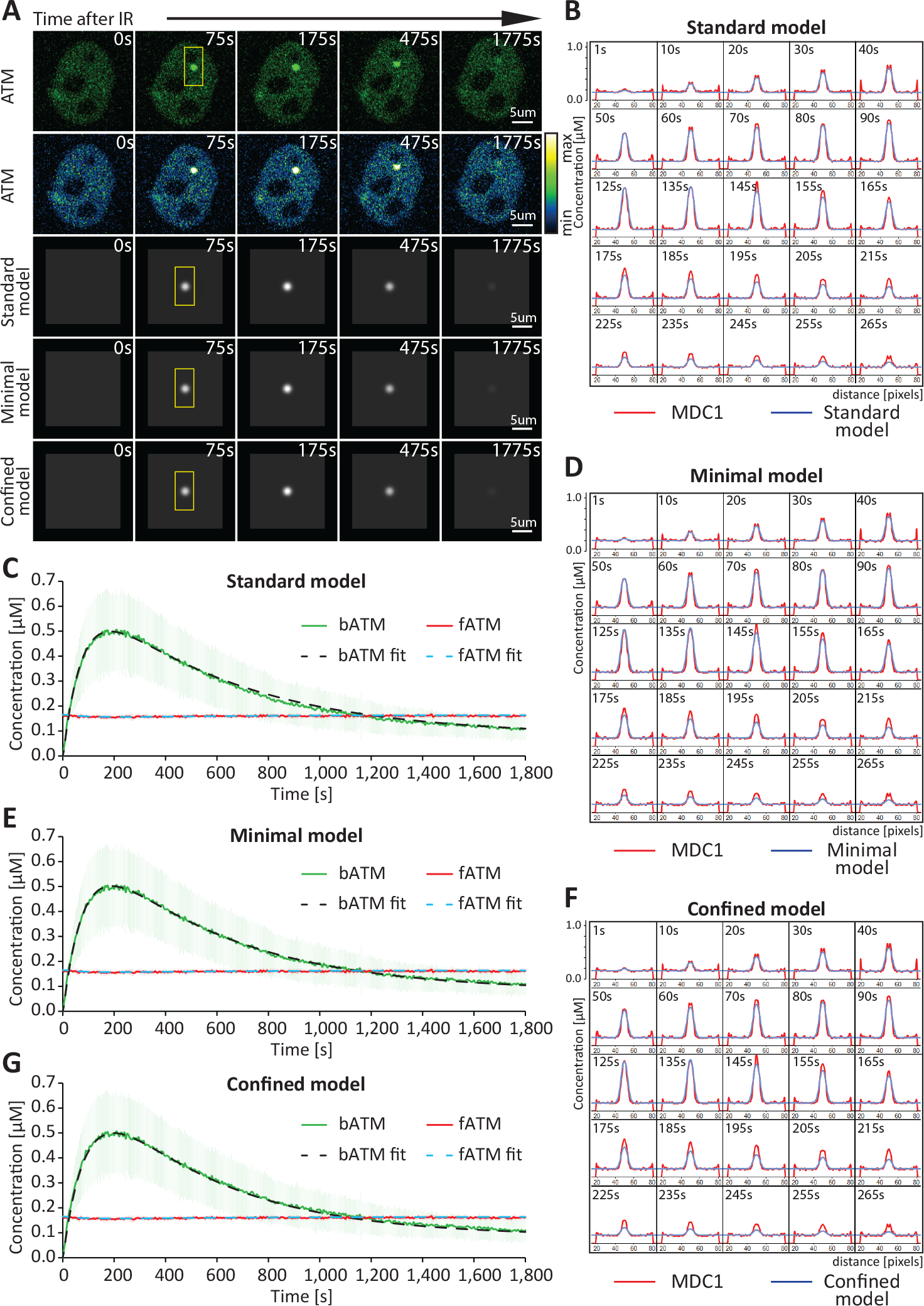
Comparison between the experimentally measured and predicted ATM dynamics at sites of complex DNA damage. (A)Comparison between time-lapse microscopy images and images generated via simulation based on the models of ATM distribution at sites of complex DNA damage. The yellow box covers a region of interest where the ATM spread presented in B, D, and F is measured. (B)Time-lapse representation of the ATM concentration profile across the micro-irradiation site (the region of interest in A). Red color indicates the measured ATM concentration spread. Blue indicates the best-fitting predicted concentration based on the standard model. (C) Graphical comparison between the experimentally measured ATM dynamics at sites of complex DNA damage and those predicted based on the standard model. Error bars represent the standard deviation. bATM, measured concentration (μM) of ATM within the DNA damage site; fATM, measured concentration (μM) of ATM outside of the damage focus; bATM fit, the best-fitting predicted concentration (μM) of ATM at the damage site, based on the standard model; fATM fit, the best-fitting predicted concentration (μM) of ATM outside of the damage site, based on the standard model. (D) Time-lapse representation of the ATM concentration profile across the micro-irradiation site (the region of interest in A). Red color indicates the measured ATM concentration spread. Blue indicates the best-fitting predicted concentration based on the minimal model. (E) Graphical comparison between the experimentally measured ATM dynamics at sites of complex DNA damage and those predicted based on the minimal model. Error bars represent the standard deviation. bATM, measured concentration (μM) of ATM within the DNA damage site; fATM, measured concentration (μM) of ATM outside of the damage focus; bATM fit, the best-fitting predicted concentration (μM) of ATM at the damage site, based on the minimal model; fATM fit, the best-fitting predicted concentration (μM) of ATM outside of the damage site, based on the minimal model. (F) Time-lapse representation of the ATM concentration profile across the micro-irradiation site (the region of interest in A). Red color indicates the measured ATM concentration spread. Blue indicates the best-fitting predicted concentration based on the confined model. (G) Graphical comparison between the experimentally measured ATM dynamics at sites of complex DNA damage and those predicted based on the confined model. Error bars represent the standard deviation. bATM, measured concentration (μM) of ATM within the DNA damage site; fATM, measured concentration (μM) of ATM outside of the damage focus; bATM fit, the best-fitting predicted concentration (μM) of ATM at the damage site, based on the confined model; fATM fit, the best-fitting predicted concentration (μM) of ATM outside of the damage site, based on the confined model.

**Figure 6.**
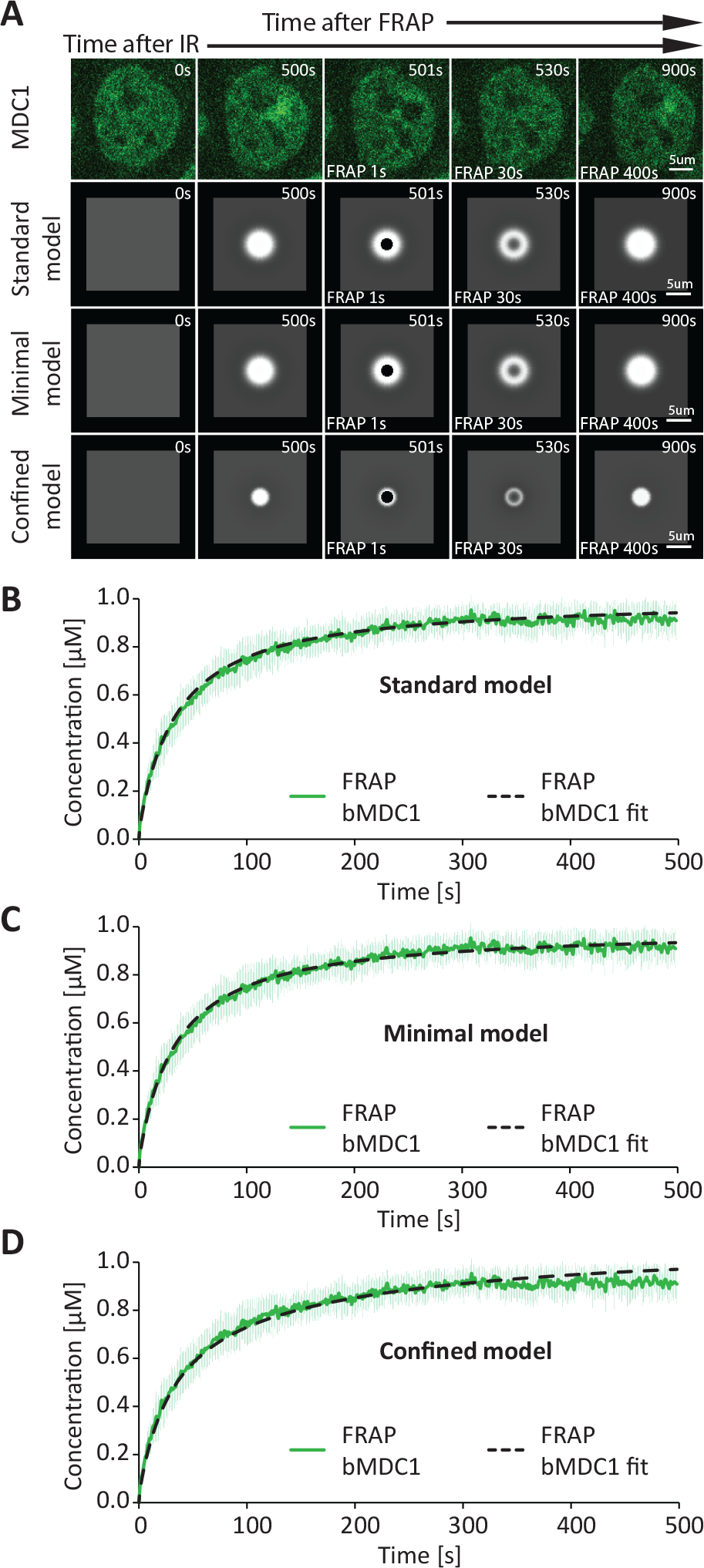
Comparison between the experimentally measured and predicted MDC1 dynamics at sites of complex DNA damage after FRAP. (A) Comparison between time-lapse microscopy images and images generated via simulation based on the models of MDC1 distribution at sites of complex DNA damage after FRAP. (B) Graphical comparison between the experimentally measured dynamic MDC1 concentration (μM) at sites of complex DNA damage after FRAP and that predicted based on the standard model. Error bars represent the standard deviation. FRAP bMDC1, measured concentration (μM) of MDC1 within the DNA damage site; FRAP bMDC1 fit, the best-fitting predicted concentration (μM) of MDC1 at the damage site, based on the standard model. (C) Graphical comparison between the experimentally measured dynamic MDC1 concentration (μM) at sites of complex DNA damage after FRAP and that predicted based on the minimal model. Error bars represent the standard deviation. FRAP bMDC1, measured concentration (μM) of MDC1 within the DNA damage site; FRAP bMDC1 fit, the best-fitting predicted concentration (μM) of MDC1 at the damage site, based on the minimal model. (D) Graphical comparison between the experimentally measured dynamic MDC1 concentration at sites of complex DNA damage after FRAP and that predicted based on the confined model. Error bars represent the standard deviation. FRAP bMDC1, measured concentration (μM) of MDC1 within the DNA damage site; FRAP bMDC1 fit, the best-fitting predicted concentration (μM) of MDC1 at the damage site, based on the confined model.

**Figure 7.**
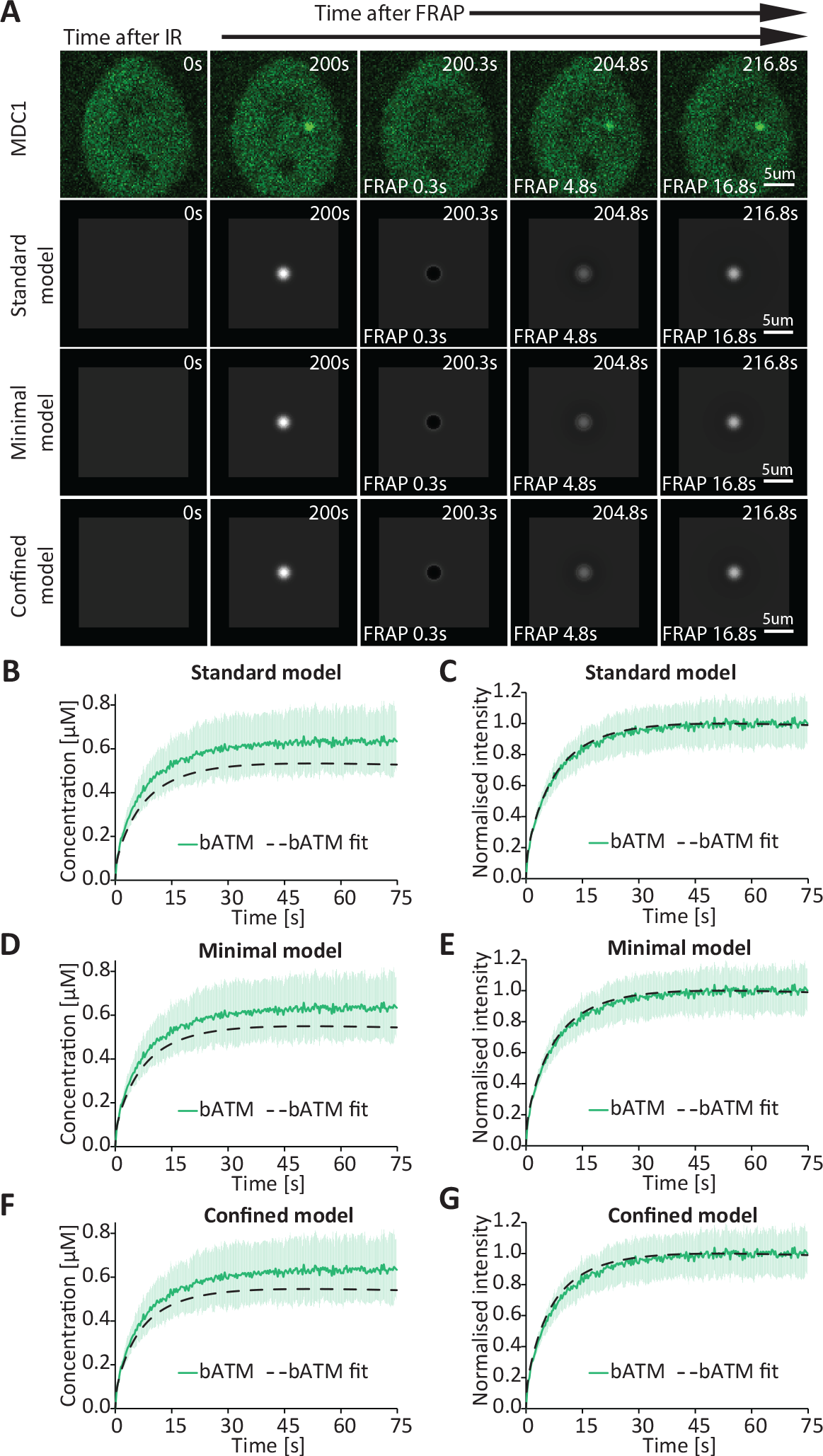
Comparison between the experimentally measured and predicted MDC1 dynamics at sites of complex DNA damage after FRAP. (A) Comparison between time-lapse microscopy images and images generated via simulation based on the models of MDC1 distribution at sites of complex DNA damage after FRAP. (B) Graphical comparison between the experimentally measured dynamic MDC1 concentration at sites of complex DNA damage after FRAP and that predicted based on the standard model. Error bars represent the standard deviation. FRAP bMDC1, measured concentration (μM) of MDC1 within the DNA damage site; FRAP bMDC1 fit, the best-fitting predicted concentration (μM) of MDC1 at the damage site, based on the standard model. (C) Graphical comparison between the normalized MDC1 kinetics at sites of complex DNA damage after FRAP and those predicted based on the standard model. Error bars represent the standard deviation. FRAP bMDC1, measured concentration (μM) of MDC1 within the DNA damage site; FRAP bMDC1 fit, the best-fitting predicted concentration (μM) of MDC1 at the damage site, based on the standard model. (D) Graphical comparison between the experimentally measured dynamic MDC1 concentration at sites of complex DNA damage after FRAP and that predicted based on the minimal model. Error bars represent the standard deviation. FRAP bMDC1, measured concentration (μM) of MDC1 within the DNA damage site; FRAP bMDC1 fit, the best-fitting predicted concentration (μM) of MDC1 at the damage site, based on the minimal model. (E) Graphical comparison between the normalized MDC1 kinetics at sites of complex DNA damage after FRAP and those predicted based on the minimal model. Error bars represent the standard deviation. FRAP bMDC1, measured concentration (μM) of MDC1 within the DNA damage site; FRAP bMDC1 fit, the best-fitting predicted concentration (μM) of MDC1 at the damage site, based on the minimal model. (F) Graphical comparison between the experimentally measured dynamic MDC1 concentration at sites of complex DNA damage after FRAP and those predicted based on the confined model. Error bars represent the standard deviation. FRAP bMDC1, measured concentration (μM) of MDC1 within the DNA damage site; FRAP bMDC1 fit, the best-fitting predicted concentration (μM) of MDC1 at the damage site, based on the confined model. (G) Graphical comparison between the normalized MDC1 kinetics at sites of complex DNA damage after FRAP and those predicted based on the confined model. Error bars represent the standard deviation. FRAP bMDC1, measured concentration (μM) of MDC1 within the DNA damage site; FRAP bMDC1 fit, the best-fitting predicted concentration (μM) of MDC1 at the damage site, based on the confined model.

We were able to find a set of parameters (Table S1) that fit the numerical solutions of the reaction-diffusion equations to the experimental data (see methods). We provide graphical representations of the proposed numerical solutions for MDC1 (Figure 4C) and ATM (Figure 5C) recruitment as well as for their depletion in the nuclear region away from the damage site. The standard model (1-9) accurately fit the measured MDC1 (Figure 4C) and ATM (Figure 5C) recruitment kinetics and quantity at the site of DNA damage as well as the simultaneous depletion of the given protein in a nuclear region away from the damage site. Furthermore, the model accurately predicted the spatiotemporal distribution of ATM (Figure 5B; Figure S3; Movie S5) and MDC1 (Figure 4B; Figure S2; Movie S4), in particular the spread of the latter away from the damage site (Figure S4 and Movie S6). The model also accurately fits the measured MDC1 (Figure 6B) and ATM (Figure 7B and 7C) kinetics of FRAP at the site of DNA damage.

A particularly impressive feature of such modeling is that it can recapitulate a scenario in which the resulting kinetics is a combination of two separate, simultaneous processes: the removal of ATM from the damage site and the ATM FRAP (Figure S5).

From fitting the model to the data, we found that the αATM effective diffusion coefficient (average D=0.059 μm^2^/s) should be 10 times smaller than the diffusion coefficient of the non-activated ATM (Table S1). A possible reason for the slow diffusion of αATM is that its transport involves binding to and unbinding from the H2AX anchored within chromatin as well as its free diffusion in between these events. The magnitude of the effective diffusion coefficient is similar to the diffusion coefficient of MDC1. The delay in MDC1 depletion from outside of the DNA damage shown in Figure 4C and the area to which MDC1 spreads are very sensitive to the value of MDC1 and αATM diffusion coefficients, which highlights the importance of their diffusion for the MDC1 spread kinetics. The standard model also implies that the concentration of lesion-bound and freely diffusing αATM are two orders of magnitude lower than that of total bound inactive ATM at the damage site (Figure S6; Movie S7).

### Minimal model explaining MDC1 spread based on diffusing activated ATM

Interestingly, the goodness of fit was not affected by changes in k_4_ (Table S1), a rate constant describing the dissociation of αATM from the damage site (Eq. 4). The MDC1 spread could therefore be explained without the presence of bound αATM at the site of damage. Rather, this model postulates that ATM is immediately released upon its activation. We will call this model the “Minimal αATM diffusion model”, or the “Minimal model”, for simplicity. Here, reactions (3) and (4) from the first model are replaced by (16). The best Minimal model fit of the experimental MDC1 (Figure 4A, E, and D; Figure S2; Movie S4) and ATM (Figure 5A, E, and D; Figure S3; Movie S5) data was as accurate as that of the Standard model, including for MDC1/ATM spatial distribution (Figure S4 and Movie S6) and MDC1/ATM FRAP data (Figure 6C; Figure 7D and 7F).

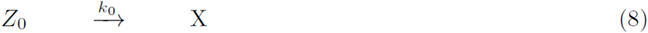

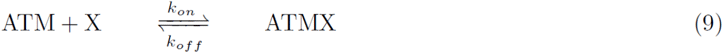

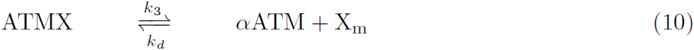

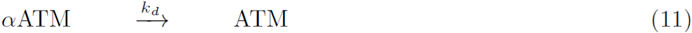

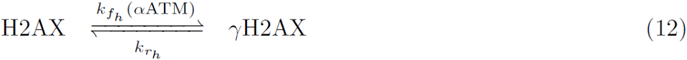

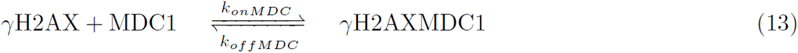

### MDC1 spread cannot be explained without the diffusion of activated ATM

We also checked whether the data can be fitted to a model in which ATM is only active when bound to the damage site. We will call this model the “Confined αATM model”, or the “Confined model”, for simplicity. The Confined model is described by the following reactions:

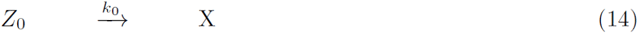

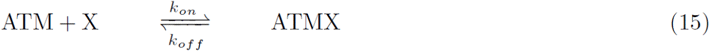

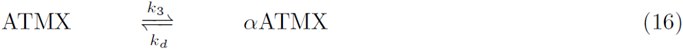

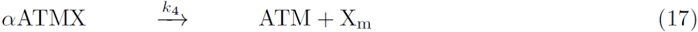

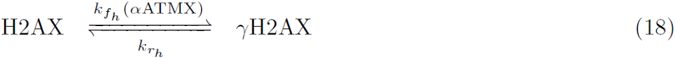

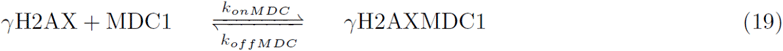

where reactions (3) and (4) from the original model are replaced by (11).

While this model fits the recruitment kinetics of ATM (Figure 5A, 5G and 5F; Figure S3; Movie S5) at the DNA damage site well, it did not fit the early steps of MDC1 recruitment to the damage site nor its delayed depletion outside of the damage site (Figure 4G). Most importantly, the model could not recapitulate MDC1 spread beyond the micro-IR-induced complex lesion (Figure 4A and 4F; Figure S2; Movie S4 and Movie S6. These observations suggest that αATM diffusion is critical for explaining MDC1 behavior during repair of complex DNA lesions.

Altogether, our results demonstrate that late recruitment of cohesin in comparison with MDC1 do not support the hypothesis that damage-induced spread of γH2AX is a result of cohesion dependent loop extrusion, at least during the repair of complex DNA damages. To explore the reason behind the γH2AX /MDC1 spread we introduce a mathematical theory that enables the spatiotemporal modelling of heterogenous chemical reactions in the cell context. Through this mathematical description, we were able to demonstrate that the damage-induced spread of γH2AX and MDC1 can be explained by simple diffusion of activated ATM.

## Discussion

In the present work, to study the nature of γH2AX/MDC1 spread at complex DNA lesions, we followed the spatiotemporal dynamics of fluorescently-tagged ATM, MDC1, NIPBL, and RAD21 through live-cell microscopy. Our measurements show that MDC1 transport in the absence of DNA damage is approximately an order of magnitude slower than expected from the pure diffusion of a similar-size protein, which is in line with a recent report demonstrating that most MDC1 molecules are indeed chromatin-bound in such conditions. After irradiation, MDC1 was quickly recruited to the damage site and subsequently spread around the DNA lesion, being heavily depleted from other parts of the nucleus. The slow MDC1 transport and its limited concentration limit foci spread after sequential irradiation at two regions of interest. In contrast to MDC1, ATM is recruited at the site of DNA damage without spreading and exhibits a fast exchange rate, close to that of freely diffusing ATM.

We developed three quantitative models of ATM and MDC1 kinetics, two of which accurately fitted our imaging data. Critical for a successful fit was the inclusion of the diffusion of αATM outside of the damage site. In the simpler of these two models, ATM dissociates from the damage site upon its activation. While diffusing, it phosphorylates H2AX histones, and the spatial extent of the γH2AX and MDC1 spread is determined by the interplay between αATM effective diffusion and the rate of its deactivation. While the former leads to phosphorylation away from the focus, the distance to which phosphorylation extends is limited by the deactivation rate.

Our model of γH2AX/MDC1 spread based on αATM 3D diffusion explains a number of measurements and observations. First, it describes the spread rate and spatial concentration profile of MDC1, as well as its exchange and recruitment kinetics following UV laser micro-irradiation. Second, our model accurately predicts slow MDC1 transport and exchange rate in the absence of DNA lesions. Third, the model faithfully describes MDC1 depletion kinetics at a region far from the lesion. Fourth, the model explains ATM exchange and recruitment kinetics, also allowing us to quantitatively predict the amount of ATM recruited at the DNA lesion. Our model is consistent with the data only if the diffusion rate of αATM is an order of magnitude slower than that of non-activated ATM (as was measured in our experiments) and similar to the MDC1 effective diffusion rate. A logical contributor to the slow diffusion rate of the activated ATM is its binding and unbinding from H2AX during phosphorylation. ATM activation is a critical step for γH2AX and MDC1 spread. It has been shown that ATM and DNA-PKs can phosphorylate H2AX, however only ATM promotes the spread of phosphorylation to large distances and high densities ^23^. ATM binds to the MRN complex via the C-terminal domain of NBS1 ^24^. It has been proposed that ATM auto-phosphorylation (at S367, S1893, S1981, S2996, and other potential sites) and/or disassociation of the ATM dimer triggers its activation ^25-27^. The loss of auto-phosphorylation sites in human ATM reduces activation ^25,28^, while mouse ATM with a mutation in the S1987 auto-phosphorylation site exhibits normal activation ^29-31^. Further, structural studies have shown that ATM activation can occur without monomerization of the ATM dimer ^26^. The αATM diffusion-based model of γH2AX/MDC1 spread proposed herein is consistent with the current mechanistic understanding of ATM activation.

It has been suggested that chromatin architecture plays an important role in the γH2AX spread, as topologically associated domains often coincide with phosphorylated H2AX regions. Based on this finding, it was proposed that chromatin contacts near DSBs correlate with γH2AX distribution, which can be explained by the 3D diffusion of αATM. In addition, it was shown that cohesin is recruited to the vicinity of a DSB, where it carries out loop extrusion, thus reshaping chromatin topology. Locally activated ATM anchored at the loop boundary was also proposed to extensively phosphorylate H2AX by modifying the DNA that is actively extruded by cohesin in a unidirectional manner at the DSB. However, cohesin depletion leads to a reduction in H2AX phosphorylation of only 10%. We show that cohesin loader NIPBL, which is necessary for cohesin recruitment at the DSBs, accumulates at a significantly slower rate than MDC1 recruitment and spread at micro-IR-induced complex DNA lesions. Cohesin subunit RAD21 exhibited recruitment kinetics in line with that of NIPBL, once again considerably slower than of MDC1. These results suggest that loop extrusion alone cannot explain the γH2AX/MDC1 spread kinetics at complex DNA lesions. The fast rate of ATM exchange, which we measured at the DNA damage site, is also hard to reconcile with continuous loop extrusion-driven H2AX phosphorylation at the DSB. In contrast, our 3D model does recapitulate the experimental data. A number of differences in experimental design may give rise to divergent results that are consistent with distinct models. First is the nature of damage. Repetitive endonuclease-mediated cutting results in persistent DSB induction. While this approach enables the specific study of a single DSB, constant cutting at the same locus does not allow one to accurately follow the sequence of repair events through time. In this manner, one cannot conclusively determine whether loop extrusion is a cause or consequence of γH2AX/MDC1 spread. Meanwhile, micro-IR induces complex DNA lesions, i.e. chromatin at the damaged region incurs a variety of damage types, including single-strand breaks, DSBs, and base modifications ^1^. Such complex lesions require the convergence and coordination of multiple repair pathways. Further, the multitude of insults may additionally alter chromatin topology. An advantage of this methodology is that damage induction occurs at a pre-defined timepoint and a confined nuclear region, which enables us to precisely follow sequential repair events. The over 200-second difference in recruitment halftime between MDC1 and RAD21 is clearly discernible through micro-IR combined with time-lapse live-cell microscopy. Such a difference cannot be detected through the sequencing-based study of repetitive DSB induction. Despite the distinct nature of complex DNA lesions relative to a single DSB, we still observe a clear spread of MDC1 at micro-IR regions. This indicates that the spread occurs, regardless of potential topological alterations in chromatin. We demonstrate that this spread cannot be explained by cohesin-mediated loop extrusion. If loop extrusion does take place at the complex DNA lesion, this occurs at a much later point in time, as indicated by the kinetics of NIPBL and RAD21. Our measurements and mathematical models indicate that the spread can be easily explained by a 3D diffusion model of αATM. This model correctly predicts the measured recruitment kinetics, exchange rate, and diffusion of aATM and MDC1 as well as the MDC1 spread at complex DNA lesions.

In summary, we employed the introduced mathematical models to describe a central step of the cellular response to DNA damage. We are confident that our mathematical theory can be applied in the study spatiotemporal fluctuations in protein concentrations in order to shed light on the molecular mechanisms underlying cellular processes.

## Materials and methods

### Cell lines

We used HeLa Kyoto cell lines stably expressing ATM-NFLAP, MDC1-LAP, and mNIPBL-NFLAP fluorescently-tagged proteins ^1,14^. The HCT-116-RAD21-mAC cell line, in which both RAD21 alleles are tagged with auxin-inducible degrons and an mClover reporter, was a kind gift from Dr. Masato Kanemaki ^21^. All cell lines were cultured in Dulbecco’s Modified Eagle Medium (DMEM) supplemented with 10% fetal bovine serum (FBS), 100 units/ml penicillin, and 100 μg/ml streptomycin at 37°C, 5% CO2. For micro-irradiation (IR) experiments and time-lapse imaging, we plated cells in MatTek glass bottom dishes (∼20% confluence) and cultured them for 48 h. Prior to image acquisition, the medium was changed to FluoroBrite DMEM medium (Thermo Fisher Scientific), containing 10% FBS and 2 mM GlutaMAX Supplement (Thermo Fisher Scientific).

### Calculation of ATM1 and MDC1 nuclear concentration

The nuclear concentrations of MDC1 and ATM were calculated as follows. We consider that MDC1 and ATM1 copy numbers per cell was 45853 and 25928 respectively as was estimated in (Hein et al., 2015). Considering that ATM and MDC1 are localized into the nucleus and that average volume of HeLa Kyoto nucleus is 0.248 pl we estimated that the concentrations of MDC1 and ATM into the HeLa Kyoto nucleus are 0.3028 μM and 0.155 μM respectively.

### Micro-irradiation (IR) and image acquisition

We performed micro-IR with an Andor Micropoint system, consisting of a 365nm dye laser pumped with a 337nm nitrogen laser at 3.5ns pulses with a pulse energy of 150 μJ. Micro-IR consisted of 10 pulses, with the 365nm dye laser output attenuated to 70% of the maximum. Cells were kept at 37°C and 5% CO2 during micro-IR experiments and image acquisition on an Andor Dragonfly 500 system equipped with a Nikon Eclipse Ti-E inverted microscope and a Nikon Perfect Focus System (PFS). We used a Nikon CFI Plan Apo 60x (NA 1.4) objective and an iXon888 EMCCD camera. Cells were imaged in three Z planes separated by 0.5 μm, at intervals of 0.5 to 5 s.

### Fluorescence Recovery After Photobleaching (FRAP)

To determine the exchange rate of DNA repair proteins at damage sites, we performed FRAP of foci after micro-IR. FRAP experiments were performed on an Andor Revolution System using a Nikon CFI Plan Apo VC 60x (NA 1.2) objective and an iXon897 EMCCD camera. After the concentration of proteins of interest in the irradiated cells reached a plateau, the micro-irradiated site was bleached (FRAP settings: 60μs dwell time, 20 repeats, 6% of the total energy of the 488nm laser -50mW).

### Micro-irradiation image and analysis

To determine the kinetics of protein recruitment after micro-IR, we performed image analysis using our CellTool software^32^, as previously described. Three Z planes for every time point were combined using a maximum intensity projection, and the images from all time points were registered to compensate for cell movement. The average intensity (I) of both the region of protein recruitment and a nearby region were measured in order to calculate the difference between intensities of these two regions. Of note, IR-induced bleaching is the same at the site of DNA damage and the nearby region at time 0, showing that the bleached region is larger than the region of protein recruitment.

Therefore, the difference between the average intensities of the region of protein recruitment and the nearby region provides the average intensity of recruited proteins, which is compensated for bleaching during UV irradiation and image acquisition. This difference was multiplied by the protein recruitment area. Thereafter, the intensity of the post-IR frame was subtracted from the obtained intensity for every time point. Using these calculations, we obtained the total intensity of recruited proteins at DNA damage sites for a given time point.

To measure the spatial distribution of tagged proteins after micro-IR, the images were analyzed as follows. A region of interest encompassing the MDC1 signal spread around the damage site was cropped^33,34^. The cropped 2D images were converted to 1D via maximum intensity projection. The center of the focus was determined automatically as the pixel with the highest intensity, and, based on this, all the images were aligned in space. MDC1 does not homogenously spread outward of the damage focus due to the heterogenous distribution of chromatin. Thus, we take the longest stretch of MDC1 spread in a given direction radiating from the focus. We are then able to follow the intensity of pixels away from the site of damage (i.e. pixels within the measured stretch) over time. To compare the observed spread to that predicted by different models, we mirror the intensity over time data, as if an analogous spread were happening in the opposite direction away from the focus.

### Image analysis of FRAP and calculation of diffusion coefficients

Image analysis of FRAP and calculation of diffusion coefficients was performed using CellTool. The average intensity in the photobleached region, in the whole cell nucleus, and in the background outside of the cell nucleus was measured. Those three values were used in order to normalize the data for acquisition bleach correction by applying the following formulas ^35^:

Double normalization:

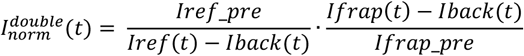

Where:

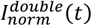– double-normalized intensity;

*Ifrap*(*t*) – measured average intensity inside the bleached spot;

*Iref*(*t*) – measured average reference or whole studied compartment (cell, nucleus, etc.) intensity;

*Iback*(*t*) – measured average background intensity outside the cell;

Subscript *_pre* means the averaging of intensity in the corresponding region of interest before bleach moment after subtraction of background intensity;

Full-scale normalization:

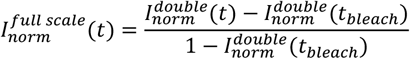

Where:

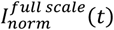 - full-scale normalized intensity;

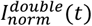 - double-normalized intensity;

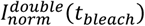 - double-normalized intensity at the time of the bleach;

The following fit of the diffusion model for an oval region of interest to the acquired normalized FRAP data was performed as described in the CellTool manual. The used model can be briefly summarized by the following formula [Soumpasis]:

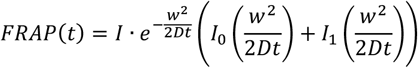

Where:

*I*_0_(*x*), *I*_1_(*x*), -modified Bessel functions.

*I* – normalizing coefficient to account the incomplete recovery;

*D* - diffusion coefficient [μm^2^s^-1^];

*w* - the radius of bleach spot [μm];

*t* – time after the bleaching [s];

The diffusion coefficient of the free protein is obtained from the FRAP of the cells that were not micro-irradiated. The diffusion coefficient of the proteins at the binding site is calculated from the analysis of the FRAP of micro-irradiated cells.

## Mathematical modeling

### Reaction-diffusion equations

The reaction-diffusion equations for N concentrations A_i_, Bi, and C_i_ are:

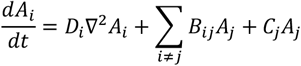

They can be treated as coupled initial-value problems over a grid of points covering the sample volume. For the case at hand, the focus site has cylindrical symmetry, so the grid consists of only a set of radial coordinates. At each time step, the diffusion term is evaluated across the grid, with the results combined with the B and C terms in the equation. To maintain stability of the diffusion terms, the time step Dt and the grid spacing Dr must satisfy the following constraint:

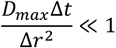

where D_max_ is the largest diffusion constant.

The stability and the convergence of the Laplacian were studied using both second- and fourth-order Laplacian finite difference expressions. The stability and the convergence of the time dependence were studied using both a first-order Euler and a fourth-order Runge-Kutta method. Using the relevant parameters for our models, Euler integration with second-order Laplacians were sufficient to provide stable, convergent results. The Laplacians were evaluated in parallel at each time step allowing the computation to be optimized on multi-core CPUs. The typical time step in the numerical solutions was 1 ms, and the grid spacing was 1 nm. The typical time of the numerical computations was around 1s.

The Nelder-Mead solver algorithm was used for automatic variation of the parameters of the models during fitting to the measured data.

We assume that the focus is in the center of a thick disc of radius R, and the concentration is symmetric in all directions, and depends only on the radius r to the center. The discrete Laplacian is calculated via

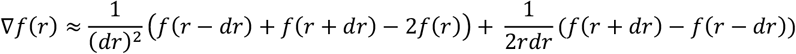

Implementation of the reaction-diffusion equations for the models is presented in supplementary information.

### Image generation from the numerical solutions

The 2D images of the model were calculated based on the 1D model arrays. The distance of every pixel of the 2D image to the center of the image was calculated, and the corresponding intensity from the 1D model array was used. Additional normalization was applied to transform the intensity of the pixels to 8-bit integer values. The maximum calculated pixel intensity during the time-lapse was transformed to the maximum value of an 8-bit integer, and all other pixels were scaled with the same factor.

## Acknowledgments

We would like to thank Petar-Bogomil Kanev for carefully reading the manuscript and his useful suggestions. HCT-116-RAD21-mAC cell line was a kind gift from Dr. Masato Kanemaki. G.D. and S.S. S acknowledge the support from the Bulgarian NSF grant# КP-06-N21-9. K.B. B. was supported by the National Science Foundation, while working at the Foundation. Any opinion, finding and conclusions or recommendations expressed in this material are those of the authors and do not necessarily reflect the views of the National Science Foundation. G. P. was partially supported by the National Science Foundation. The authors acknowledge the support from the Sofia Euro-Bioimaging node of NRIR.

## Author contributions

S.S.S, designed the experiments, G.D. and S.S.S. conducted the experiments. G.D., G.P., B.K., G.G., K.B.B., and S.S.S developed the mathematical model, G.D. and B.K. wrote the numerical solution software. S.S.S., K.B.B., and G.P. wrote the paper. All authors commented on and edited the text.

## Declaration of interests

The authors declare no competing interests.

## Notes

### Competing Interest Statement

The authors have declared no competing interest.

