## Supplemental information for "Diffusion of activated ATM explains γH2AX and MDC1 spread beyond the DNA damage site"

#### 1. Reaction-diffusion models

The damage site recruits free ATM protein, which binds to the damage site and is activated. The activated ATM then facilitates the phosphorilation of histones to which MDC1 binds. Here we consider 4 different reaction-diffusion models which could describe this system.

Throughout we make the following geometric assumption: The cell is cylindrical with radius $R$ and the damage site, which we refer to as "focus", is in the middle. This geometry enables us to use symmetry and reduce dimensionality.

We will consider the interaction between the following species: the DNA damage (denoted by X, which are treated as parameters), the enzyme ATM (denoted by ATM) and the activated ATM (denoted by $\alpha$ATM), the histones (denoted by H2AX) and phosphirlated histones (denoted by $\gamma$H2AX). The free MDC1 is denoted by $S$ and the bound MDC1 is denoted by $\gamma$H2AXMDC1 for the two foci. The spatial dependence is implicit everywhere. X has different initial values at the two foci, and is 0 everywhere else in the nucleus. The histones are present everywhere. The species which diffuse are the ATM and activated ATM and the free MDC1. The diffusion constants for ATM is $D_a$, for activated ATM it is $D_p$, and for free MDC1 is $D_m$.

##### 1.1. Standard model.

Activation of ATM from ATM to $\alpha$ATM happens only in the damaged site and modifies the damage irreversibly. The damage has two forms, X and $X_1$, where X leads to the complex ATMX and $X_1$ leads to the bound state $B$. The reactions can be reversible, and the ATM–damage complex can transition into three states.

| | | |
|---|---|---|
| $Z_0 \xrightarrow{k_0}$ | X | damage to substrate |
| $\mathrm{ATM} + \mathrm{X} \underset{k_{off}}{\overset{k_{on}}{\rightleftharpoons}}$ | ATMX | ATM forms a complex in the damaged site, |
| $\mathrm{ATMX} \underset{k_d}{\overset{k_3}{\rightleftharpoons}}$ | $\alpha$ATMX | activation of ATM to $\alpha$ATMX complex |
| $\alpha\mathrm{ATMX} \xrightarrow{\mathrm{k_4}}$ | $\alpha\mathrm{ATM} + \mathrm{X_m}$ | activated ATM and modified damage site |
| $Z_1 \xrightarrow{k_0}$ | $X_1$ | damage to substrate |
| $\mathrm{ATM} + \mathrm{X_1} \xrightarrow{\mathrm{k_1}}$ | $B$ | some of ATM remains bound to the focus |
| $\alpha\mathrm{ATM} \xrightarrow{\mathrm{k_d}}$ | ATM | deactivation of ATM |

Following the dynamics of ATM, the activated ATM phosphorilates the histones H2AX to $\gamma$H2AX. Once the histones are phosphorilated by the activated ATM, MDC1 then attaches to them. In the main model both $\alpha$ATM and $\alpha$ATMX phosphorilate the histone at the given location and act as enzymes.

$$\mathrm{H2AX} \xrightleftharpoons[k_{r_h}]{k_{f_h}(\alpha\mathrm{ATM}+\alpha\mathrm{ATMX})} \gamma\mathrm{H2AX} \qquad \text{phosphorilation of histones by ATM}$$

$$\gamma\mathrm{H2AX} + \mathrm{MDC1} \xrightleftharpoons[k_{offMDC}]{k_{onMDC}} \gamma\mathrm{H2AXMDC1} \qquad \text{free MDC1 binds to phosphorilated histones}$$

This model translates to the following differential equations, where $\Delta = \nabla^2$ is the Laplace operator. All concentrations here are functions $\mathbb{R}^{3+1} \to \mathbb{R}_{\geq 0}$.

$$\frac{d\ \mathrm{ATM}}{dt} = -k_{on}\mathrm{ATM} * \mathrm{X} - \mathrm{k_1 ATM}\Bigg(\mathrm{Z_1(t=0)(1-e^{-k_0 t}) - B}\Bigg) \tag{1}$$
$$+ k_{off}\mathrm{ATMX} + \mathrm{k_d}\alpha\mathrm{ATM} + \mathrm{D_a}\Delta\mathrm{ATM}$$
$$\frac{d\ \mathrm{X}}{dt} = -k_{on}\mathrm{ATM} * \mathrm{X} + \mathrm{k_{off}ATMX} + \mathrm{k_0 Z_0 e^{-k_0 * t}} \tag{2}$$
$$\frac{d\ \mathrm{ATMX}}{dt} = k_{on}\mathrm{ATM} * \mathrm{X} - (\mathrm{k_{off}} + \mathrm{k_3})\mathrm{ATMX} + \mathrm{k_d}\alpha\mathrm{ATMX} \tag{3}$$
$$\frac{d\ \alpha\mathrm{ATM}}{dt} = k_4\alpha\mathrm{ATMX} - \mathrm{k_d P} + \mathrm{D_p}\Delta\alpha\mathrm{ATM} \tag{4}$$
$$\frac{d\ \alpha\mathrm{ATMX}}{dt} = k_3\mathrm{ATMX} - \mathrm{k_4}\alpha\mathrm{ATMX} - \mathrm{k_d}\alpha\mathrm{ATMX} \tag{5}$$
$$\frac{d\ B}{dt} = k_1\mathrm{ATM} * \Bigg(\mathrm{Z_1(t=0)(1-e^{-k_0 t}) - B}\Bigg) \tag{6}$$

For MDC1 we have the following equations

$$\frac{d\ \mathrm{H2AX}}{dt} = -k_{f_h} * (\alpha\mathrm{ATM} + \alpha\mathrm{ATMX}) * \mathrm{H2AX} + \mathrm{k_{r_h}} * \gamma\mathrm{H2AX} \tag{7}$$
$$\frac{d\ \gamma\mathrm{H2AX}}{dt} = k_{f_h} * (\alpha\mathrm{ATM} + \alpha\mathrm{ATMX}) * \mathrm{H2AX} - \mathrm{k_{r_h}} * \gamma\mathrm{H2AX} \tag{8}$$
$$+ k_{off}\gamma\mathrm{H2AXMDC1} - \mathrm{k_{on}}\gamma\mathrm{H2AX} * \mathrm{MDC1}$$
$$\frac{d\ \gamma\mathrm{H2AXMDC1}}{dt} = -k_{offMDC} * \gamma\mathrm{H2AXMDC1} + \mathrm{k_{onMDC}}\gamma\mathrm{H2AX} * \mathrm{MDC1} \tag{9}$$
$$\frac{d\ \mathrm{MDC1}}{dt} = k_{offMDC} * \gamma\mathrm{H2AXMDC1} - \mathrm{k_{onMDC}}\gamma\mathrm{H2AX} * \mathrm{MDC1} + \mathrm{D_m}\Delta\mathrm{MDC1} \tag{10}$$

Note that here we used the fact that

$$\frac{d\ \mathrm{X_1}}{dt} + \frac{dB}{dt} = k_0 Z_1(t) = k_0 Z_1(0) e^{-k_0 t}$$

and also that $\mathrm{X_1(0) = 0, B(0) = 0}$, to solve for $\mathrm{X_1 + B = Z_1(0)(1 - e^{-k_0 t})}$ and then substitute it in the equation $\frac{dB}{dt} = k_1\mathrm{ATMX_1} = \mathrm{k_1 ATM(Z_1(0)(1-e^{-k_0 t}) - B)}$.

Some of the initial values can be determined from the data. Assuming that ATM's initial concentration (also number of molecules at the focus) is much bigger than the damage X and $\mathrm{X_1}$, at steady state each initial damage site would be bound to an ATM molecule, so $B(t=\infty) = \mathrm{X_1(t=0)}$ and $\mathrm{ATMX(t=\infty) = X(t=0)}$. Also, $\mathrm{ATM(t=0)}$ is the initial concentration of $ATM$, all of which is free inactive before the damage.

1.2. **Minimal model: no $\alpha$ATMX.** Here the intermediate compound $\alpha$ATMX is missing, i.e. there is no active $\alpha$ATM bound to the damage site:

$$
\begin{aligned}
Z_0 &\xrightarrow{k_0} & \text{X} && \text{damage to substrate}\\
\text{ATM} + \text{X} &\underset{k_{off}}{\overset{k_{on}}{\rightleftharpoons}} & \text{ATMX} && \text{ATM forms a complex in the damaged site,}\\
\text{ATMX} &\underset{k_d}{\overset{k_3}{\rightleftharpoons}} & \alpha\text{ATM} + \text{X}_\text{m} && \text{activation of ATM}\\
Z_1 &\xrightarrow{k_0} & \text{X}_1 && \text{damage to substrate}\\
\text{ATM} + \text{X}_1 &\xrightarrow{\text{k}_1} & B && \text{some of ATM remains bound to the focus}\\
\alpha\text{ATM} &\xrightarrow{\text{k}_\text{d}} & \text{ATM} && \text{deactivation of ATM}
\end{aligned}
$$

Here thus only $\alpha$ATM phosphorilates the histones.

$$
\begin{aligned}
\text{H2AX} &\underset{k_{r_h}}{\overset{k_{f_h}(\alpha\text{ATM})}{\rightleftharpoons}} \gamma\text{H2AX} \quad \text{phosphorilation of histones by ATM}\\
\gamma\text{H2AX} + \text{MDC1} &\underset{k_{offMDC}}{\overset{k_{onMDC}}{\rightleftharpoons}} \gamma\text{H2AXMDC1} \quad \text{free MDC1 binds to phosphorilated histones}
\end{aligned}
$$

The corresponding equations are as follows:

$$
\begin{aligned}
\frac{d\ \text{ATM}}{dt} &= -k_{on}\text{ATM} * \text{X} - \text{k}_1\text{ATM}\Bigg(\text{Z}_1(\text{t}=0)(1-\text{e}^{-\text{k}_0\text{t}}) - \text{B}\Bigg) && (11)\\
&\quad + k_{off}\text{ATMX} + \text{k}_\text{d}\alpha\text{ATM} + \text{D}_\text{a}\Delta\text{ATM}\\
\frac{d\ \text{X}}{dt} &= -k_{on}\text{ATM} * \text{X} + \text{k}_\text{off}\text{ATMX} + \text{k}_0\text{Z}_0\text{e}^{-\text{k}_0 * \text{t}} && (12)\\
\frac{d\ \text{ATMX}}{dt} &= k_{on}\text{ATM} * \text{X} - (\text{k}_\text{off} + \text{k}_3)\text{ATMX} + \text{k}_\text{d}\alpha\text{ATM} && (13)\\
\frac{d\ \alpha\text{ATM}}{dt} &= k_4\text{ATMX} - \text{k}_\text{d}\text{P} + \text{D}_\text{p}\Delta\alpha\text{ATM} && (14)\\
\frac{d\ B}{dt} &= k_1\text{ATM} * \Bigg(\text{Z}_1(\text{t}=0)(1-\text{e}^{-\text{k}_0\text{t}}) - \text{B}\Bigg) && (15)
\end{aligned}
$$

and the MDC1 equations are

$$
\begin{aligned}
\frac{d\ \text{H2AX}}{dt} &= -k_{f_h} * (\alpha\text{ATM}) * \text{H2AX} + \text{k}_{\text{r}_\text{h}} * \gamma\text{H2AX} && (16)\\
\frac{d\ \gamma\text{H2AX}}{dt} &= k_{f_h} * (\alpha\text{ATM}) * \text{H2AX} - \text{k}_{\text{r}_\text{h}} * \gamma\text{H2AX} && (17)\\
&\quad + k_{off}\gamma\text{H2AXMDC1} - \text{k}_\text{on}\gamma\text{H2AX} * \text{MDC1}\\
\frac{d\ \gamma\text{H2AXMDC1}}{dt} &= -k_{offMDC} * \gamma\text{H2AXMDC1} + \text{k}_\text{onMDC}\gamma\text{H2AX} * \text{MDC1} && (18)\\
\frac{d\ \text{MDC1}}{dt} &= k_{offMDC} * \gamma\text{H2AXMDC1} - \text{k}_\text{onMDC}\gamma\text{H2AX} * \text{MDC1} + \text{D}_\text{m}\Delta\text{MDC1} && (19)
\end{aligned}
$$

1.3. **Confined model: no active $\alpha$ATM.** Here ATM is only active when bound to the damage site and thus no $\alpha$ATM is formed.

$$
\begin{array}{rcll}
Z_0 \xrightarrow{k_0} & \mathrm{X} & \text{damage to substrate} \\
\mathrm{ATM} + \mathrm{X} \underset{k_{off}}{\overset{k_{on}}{\rightleftharpoons}} & \mathrm{ATMX} & \text{ATM forms a complex in the damaged site,} \\
\mathrm{ATMX} \underset{k_d}{\overset{k_3}{\rightleftharpoons}} & \alpha\mathrm{ATMX} & \text{activation of ATM} \\
\alpha\mathrm{ATMX} \xrightarrow{\mathrm{k_4}} & \mathrm{ATM} + \mathrm{X_m} & \text{deactivation of ATM} \\
Z_1 \xrightarrow{k_0} & \mathrm{X_1} & \text{damage to substrate} \\
\mathrm{ATM} + \mathrm{X_1} \xrightarrow{\mathrm{k_1}} & B & \text{some of ATM remains bound to the focus}
\end{array}
$$

Here only $\alpha$ATMX phosphorilates the histones.

$$\mathrm{H2AX} \underset{k_{r_h}}{\overset{k_{f_h}(\alpha\mathrm{ATMX})}{\rightleftharpoons}} \gamma\mathrm{H2AX} \quad \text{phosphorilation of histones by ATM} \tag{20}$$

$$\gamma\mathrm{H2AX} + \mathrm{MDC1} \underset{k_{offMDC}}{\overset{k_{onMDC}}{\rightleftharpoons}} \gamma\mathrm{H2AXMDC1} \quad \text{free MDC1 binds to phosphorilated histones} \tag{21}$$

The equations are:

$$\frac{d\ \mathrm{ATM}}{dt} = -k_{on}\mathrm{ATM} * \mathrm{X} - \mathrm{k_1ATM}\left(\mathrm{Z_1(t=0)(1-e^{-k_0t}) - B}\right) \tag{22}$$
$$+ k_{off}\mathrm{ATMX} + \mathrm{k_4}\alpha\mathrm{ATMX} + \mathrm{D_a}\Delta\mathrm{ATM}$$

$$\frac{d\ \mathrm{X}}{dt} = -k_{on}\mathrm{ATM} * \mathrm{X} + \mathrm{k_{off}ATMX} + \mathrm{k_0Z_0e^{-k_0*t}} \tag{23}$$

$$\frac{d\ \mathrm{ATMX}}{dt} = k_{on}\mathrm{ATM} * \mathrm{X} - (\mathrm{k_{off}} + \mathrm{k_3})\mathrm{ATMX} + \mathrm{k_d}\alpha\mathrm{ATMX} \tag{24}$$

$$\frac{d\ \alpha\mathrm{ATMX}}{dt} = k_3\mathrm{ATMX} - (\mathrm{k_4} + \mathrm{k_d})\alpha\mathrm{ATMX} \tag{25}$$

$$\frac{dB}{dt} = k_1\mathrm{ATM} * \left(\mathrm{Z_1(t=0)(1-e^{-k_0t}) - B}\right) \tag{26}$$

and MDC1 equations:

$$\frac{d\ \mathrm{H2AX}}{dt} = -k_{f_h} * (\alpha\mathrm{ATMX}) * \mathrm{H2AX} + \mathrm{k_{r_h}} * \gamma\mathrm{H2AX} \tag{27}$$

$$\frac{d\ \gamma\mathrm{H2AX}}{dt} = k_{f_h} * (\alpha\mathrm{ATMX}) * \mathrm{H2AX} - \mathrm{k_{r_h}} * \gamma\mathrm{H2AX} \tag{28}$$
$$+ k_{off}\gamma\mathrm{H2AXMDC1} - \mathrm{k_{on}}\gamma\mathrm{H2AX} * \mathrm{MDC1}$$

$$\frac{d\ \gamma\mathrm{H2AXMDC1}}{dt} = -k_{offMDC} * \gamma\mathrm{H2AXMDC1} + \mathrm{k_{onMDC}}\gamma\mathrm{H2AX} * \mathrm{MDC1} \tag{29}$$

$$\frac{d\ \mathrm{MDC1}}{dt} = k_{offMDC} * \gamma\mathrm{H2AXMDC1} - \mathrm{k_{onMDC}}\gamma\mathrm{H2AX} * \mathrm{MDC1} + \mathrm{D_m}\Delta\mathrm{MDC1} \tag{30}$$

#### 2. FRAP models

In the FRAP experiment, the site is bleached at a given time $T$ after the initial damage. Before that all particles are light. Particles in the bleached area become dark, and the dynamics continues. In order to model this we introduce light and dark species for each compound (each form of ATM), which we denote with subscripts $\ell$ and $d$ respectively.

At the time of the photobleaching at the focus we have the following conditions. At time $t = T$ all light species in a disc or radius $r$ near the focus become dark, so the values of the light species become 0, the dark values are equal to the concentration of the corresponding light species at $t = T-$ at the given spot. All other dark values are set to 0, and the light values remain the same.

$$\mathrm{ATM_l(t=T+) = ATM_l(t=T-), ATM_d(t=T+) = 0, \alpha ATM_l(t=T+) = \alpha ATM_l(t=T-), \alpha ATM_d(t=T+) = 0}$$

$$B_l(t=T+) = 0, B_d(t=T+) = B_l(t=T-), \mathrm{ATMX_d(t=T+) = ATMX_l(t=T-), ATMX_l(t=T+) = 0}$$

Before the FRAP experiment the dynamics evolves according to the models from Section 1. We show this here with the Standard model, the other models are implemented completely analogously.

$$\begin{array}{rcl} \mathrm{ATM + X} \underset{k_{off}}{\overset{k_{on}}{\rightleftharpoons}} & \mathrm{ATMX} & \text{ATM forms a complex in the damaged site,} \\ \mathrm{ATMX} \underset{k_d}{\overset{k_3}{\rightleftharpoons}} & \alpha\mathrm{ATMX} & \text{activation of ATM to } \alpha\text{ATMX complex} \\ \alpha\mathrm{ATMX} \xrightarrow{\mathrm{k_4}} & \alpha\mathrm{ATM + X_p} & \text{activated ATM and modified damage site} \\ \mathrm{ATM + X_1} \xrightarrow{\mathrm{k_1}} & B & \text{some of ATM remains bound to the focus} \\ \alpha\mathrm{ATM} \xrightarrow{\mathrm{k_d}} & \mathrm{ATM} & \text{deactivation of ATM} \end{array}$$

Whose equations are given by

$$\frac{d\ \mathrm{ATM}}{dt} = -k_{on}\mathrm{ATM} * \mathrm{X} - \mathrm{k_1 ATM(X_1(t=0) - B) + k_{off} ATMX + k_d \alpha ATM + D_a \Delta ATM} \tag{31}$$

$$\frac{d\ \mathrm{X}}{dt} = -k_{on}\mathrm{ATM} * \mathrm{X} + \mathrm{k_{off} ATMX} \tag{32}$$

$$\frac{d\ \mathrm{ATMX}}{dt} = k_{on}\mathrm{ATM} * \mathrm{X} - \mathrm{(k_{off} + k_3) ATMX + k_d \alpha ATMX} \tag{33}$$

$$\frac{d\ \alpha\mathrm{ATM}}{dt} = k_4 \alpha\mathrm{ATMX} - \mathrm{k_d} \alpha\mathrm{ATM} + \mathrm{D_p} \Delta\alpha\mathrm{ATM} \tag{34}$$

$$\frac{d\ \alpha\mathrm{ATMX}}{dt} = k_3\mathrm{ATMX} - \mathrm{k_4} \alpha\mathrm{ATMX} - \mathrm{k_d} \alpha\mathrm{ATMX} \tag{35}$$

$$\frac{d\ B}{dt} = k_1 \mathrm{ATM} * \mathrm{(X_1(t=0) - B)} \tag{36}$$

$$\frac{d\ \mathrm{X_1}}{dt} = -k_1 \mathrm{ATM} * \mathrm{X_1} \tag{37}$$

After photobleaching at time $t = T$ we now have light and dark species, whose dynamics is governed as follows

$$\frac{d\ \mathrm{ATM}_\ell}{dt} = -k_{on}\mathrm{ATM}_\ell * \mathrm{X} + (\mathrm{k_{off}})\mathrm{ATMX}_\ell + \mathrm{k_d}\alpha\mathrm{ATM}_\ell + \mathrm{D_a}\Delta\mathrm{ATM}_\ell \tag{38}$$
$$\frac{d\ \mathrm{ATMX}_\ell}{dt} = k_{on}\mathrm{ATM}_\ell * \mathrm{X} - (\mathrm{k_{off}} + \mathrm{k_3})\mathrm{ATMX}_\ell + \mathrm{k_d}\alpha\mathrm{ATMX}_\ell \tag{39}$$
$$\frac{d\ \alpha\mathrm{ATM}_\ell}{dt} = k_4\alpha\mathrm{ATMX}_\ell - \mathrm{k_d}\alpha\mathrm{ATM}_\ell + \mathrm{D_p}\Delta\alpha\mathrm{ATM}_\ell \tag{40}$$
$$\frac{d\ \alpha\mathrm{ATMX}_\ell}{dt} = k_3\mathrm{ATMX}_\ell - \mathrm{k_4}\alpha\mathrm{ATMX}_\ell - \mathrm{k_d}\alpha\mathrm{ATMX}_\ell \tag{41}$$
$$\frac{d\ \mathrm{ATM_d}}{dt} = -k_{on}\mathrm{ATM_d} * \mathrm{X} + (\mathrm{k_{off}})\mathrm{ATMX_d} + \mathrm{k_d}\alpha\mathrm{ATM_d} + \mathrm{D_a}\Delta\mathrm{ATM_d} \tag{42}$$
$$\frac{d\ \mathrm{ATMX_d}}{dt} = k_{on}\mathrm{ATM_d} * \mathrm{X} - (\mathrm{k_{off}} + \mathrm{k_3})\mathrm{ATMX_d} + \mathrm{k_d}\alpha\mathrm{ATMX_d} \tag{43}$$
$$\frac{d\ \alpha\mathrm{ATM_d}}{dt} = k_4\alpha\mathrm{ATMX_d} - \mathrm{k_d}\alpha\mathrm{ATM_d} + \mathrm{D_p}\Delta\alpha\mathrm{ATM_d} \tag{44}$$
$$\frac{d\ \alpha\mathrm{ATMX_d}}{dt} = k_3\mathrm{ATMX_d} - \mathrm{k_4}\alpha\mathrm{ATMX_d} - \mathrm{k_d}\alpha\mathrm{ATMX_d} \tag{45}$$
$$\frac{d\ \mathrm{X}}{dt} = -k_{on}(\mathrm{ATM}_\ell + \mathrm{ATM_d}) * \mathrm{X} + \mathrm{k_{off}}(\mathrm{ATMX}_\ell + \mathrm{ATMX_d}) \tag{46}$$

For the model with inhibitor we have the same dynamics for $t < T$.

| | | |
|---|---|---|
| $\mathrm{ATM} + \mathrm{X} \underset{k_{off}}{\overset{k_{on}}{\rightleftharpoons}}$ ATMX | | ATM forms a complex in the damaged site, |
| $\mathrm{ATMX} \xrightarrow{\mathrm{k_5}}$ | $\mathrm{ATM} + \mathrm{X_m}$ | no activation of ATM |
| $\mathrm{ATM} + \mathrm{X_1} \xrightarrow{\mathrm{k_{on}}}$ | $B$ | some of ATM remains bound to the focus |

Equations:

$$\frac{d\ \mathrm{ATM}}{dt} = -k_{on} * \mathrm{XATM} - \mathrm{k_{on}}\mathrm{ATM}(\mathrm{X_1}(\mathrm{t}=0) - \mathrm{B}) + (\mathrm{k_{off}} + \mathrm{k_5})\mathrm{ATMX} + \mathrm{D_a}\Delta\mathrm{ATM} \tag{47}$$
$$\frac{d\ \mathrm{ATMX}}{dt} = k_{on}\mathrm{XATM} - (\mathrm{k_{off}} + \mathrm{k_5})\mathrm{ATMX} \tag{48}$$
$$\frac{d\ \mathrm{X}}{dt} = -k_{on}\mathrm{XATM} + \mathrm{k_{off}}\mathrm{ATMX} \tag{49}$$
$$\frac{d\ B}{dt} = k_{on}\mathrm{ATM} * (\mathrm{X_1}(\mathrm{t}=0) - \mathrm{B}) \tag{50}$$

After photobleaching the system at the presence of the inhibitor, we are assuming that $\mathrm{X_1}$ has all been converted to $B$ and the last reaction does not happen.

$$\frac{d\ \mathrm{ATM_l}}{dt} = -k_{on} * \mathrm{XATM_l} + (\mathrm{k_{off}} + \mathrm{k_5})\mathrm{ATMX_l} + \mathrm{D_a}\Delta\mathrm{ATM_l} \tag{51}$$
$$\frac{d\ \mathrm{ATMX_l}}{dt} = k_{on}\mathrm{XATM_l} - (\mathrm{k_{off}} + \mathrm{k_5})\mathrm{ATMX_l} \tag{52}$$
$$\frac{d\ \mathrm{ATM_d}}{dt} = -k_{on} * \mathrm{XATM_d} + (\mathrm{k_{off}} + \mathrm{k_5})\mathrm{ATMX_d} + \mathrm{D_a}\Delta\mathrm{ATM_d} \tag{53}$$
$$\frac{d\ \mathrm{ATMX_d}}{dt} = k_{on}\mathrm{XATM_d} - (\mathrm{k_{off}} + \mathrm{k_5})\mathrm{ATMX_d} \tag{54}$$
$$\frac{d\ \mathrm{X}}{dt} = -k_{on}\mathrm{X}(\mathrm{ATM}_\ell + \mathrm{ATM_d}) + \mathrm{k_{off}}(\mathrm{ATMX}_\ell + \mathrm{ATMX_d}) \tag{55}$$

2.1. **MDC1 FRAP experiment.** Again, we run the original experiment with the equations above until a time $t = T$, when all MDC molecules are only light. We bleach a region of radius $r$ around the focus at time

$t = T$ at which point all MDC (bound and free) in the focus becomes dark, and the reactions continue with two types of MDC – light and dark:

Before the bleaching everything is light and we have the following reactions:

$t < T$

$$\frac{d\ \mathrm{H2AX}}{dt} = -k_{f_h} * (\alpha\mathrm{ATM} + \alpha\mathrm{ATMX}) * \mathrm{H2AX} + \mathrm{k_{r_h}} * \gamma\mathrm{H2AX}$$

$$\frac{d\ \gamma\mathrm{H2AX}}{dt} = k_{f_h} * (\alpha\mathrm{ATM} + \alpha\mathrm{ATMX}) * \mathrm{H2AX} - \mathrm{k_{r_h}} * \gamma\mathrm{H2AX} + \mathrm{k_{off}}\gamma\mathrm{H2AXMDC1} - \mathrm{k_{on}}\gamma\mathrm{H2AX} * \mathrm{MDC1}$$

$$\frac{d\ \gamma\mathrm{H2AXMDC1}}{dt} = -k_{off} * \gamma\mathrm{H2AXMDC1} + \mathrm{k_{on}}\gamma\mathrm{H2AX} * \mathrm{MDC1}$$

$$\frac{d\ \mathrm{MDC1}}{dt} = k_{off} * \gamma\mathrm{H2AXMDC1} - \mathrm{k_{on}}\gamma\mathrm{H2AX} * \mathrm{MDC1} + \mathrm{D_m}\Delta\mathrm{MDC1}$$

At time $t = T$ we set for $||x - focus|| > r$:

$\mathrm{MDC1}_\ell(\mathrm{x}) = \mathrm{MDC1}(\mathrm{x})$ , $\gamma\mathrm{H2AXMDC1}_\ell(\mathrm{x}) = \gamma\mathrm{H2AXMDC1}(\mathrm{x})$, $\gamma\mathrm{H2AXMDC1_d}(\mathrm{x}) = 0$, $\mathrm{MDC1_d}(\mathrm{x}) = 0$.

For $||x - focus|| \leq r$:

$\mathrm{MDC1}_\ell(\mathrm{x}) = 0$, $\gamma\mathrm{H2AXMDC1}_\ell(\mathrm{x}) = 0$, $\mathrm{MDC1_d}(\mathrm{x}) = \mathrm{MDC1}(\mathrm{x})$, $\gamma\mathrm{H2AXMDC1_d}(\mathrm{x}) = \gamma\mathrm{H2AXMDC1}(\mathrm{x})$.

And the system of equations becomes:

$t \geq T$

$$\frac{d\ \mathrm{H2AX}}{dt} = -k_{f_h} * (\alpha\mathrm{ATM} + \alpha\mathrm{ATMX}) * \mathrm{H2AX} + \mathrm{k_{r_h}} * \gamma\mathrm{H2AX}$$

$$\begin{aligned}\frac{d\ \gamma\mathrm{H2AX}}{dt} &= k_{f_h} * (\alpha\mathrm{ATM} + \alpha\mathrm{ATMX}) * \mathrm{H2AX} - \mathrm{k_{r_h}} * \gamma\mathrm{H2AX} \\ &\quad + k_{off}(\gamma\mathrm{H2AXMDC1}_\ell + \gamma\mathrm{H2AXMDC1_d}) - \mathrm{k_{on}}\gamma\mathrm{H2AX} * (\mathrm{MDC1}_\ell + \mathrm{MDC1_d})\end{aligned}$$

$$\frac{d\ \gamma\mathrm{H2AXMDC1}_\ell}{dt} = -k_{off} * \gamma\mathrm{H2AXMDC1}_\ell + \mathrm{k_{on}}\gamma\mathrm{H2AX} * \mathrm{MDC1}_\ell$$

$$\frac{d\ \mathrm{MDC1}_\ell}{dt} = k_{off} * \gamma\mathrm{H2AXMDC1}_\ell - \mathrm{k_{on}}\gamma\mathrm{H2AX} * \mathrm{MDC1}_\ell + \mathrm{D_m}\Delta\mathrm{MDC1}_\ell$$

$$\frac{d\ \gamma\mathrm{H2AXMDC1_d}}{dt} = -k_{off} * \gamma\mathrm{H2AXMDC1_d} + \mathrm{k_{on}}\gamma\mathrm{H2AX} * \mathrm{MDC1_d}$$

$$\frac{d\ \mathrm{MDC1_d}}{dt} = k_{off} * \gamma\mathrm{H2AXMDC1_d} - \mathrm{k_{on}}\gamma\mathrm{H2AX} * \mathrm{MDC1_d} + \mathrm{D_m}\Delta\mathrm{MDC1_d}$$

2.2. **Timescale change.** In order to implement and fit all models and experiments together with the same constants, we need to use the same timescale and $\delta t$. This is done as follows in general. Let the first model's experiment be on a time range from $t = 0$ to $t = T_1$, and the second model run on time range from $t = 0$ to $t = T_2$. Assuming we have equal number of time-point measurements (i.e. both arrays have equal lengths), we can run the models on the same time scale 0 to $T_1$ by multiplying the second model's equations by

$$\tau = \frac{T_2}{T_1}.$$

If the models are $\frac{d\vec{F}_i(t)}{dt} = \vec{G}_i(\vec{F}_i(t))$ then the two can put together as:

$$\frac{d\ \vec{F}_1(t)}{dt} = \vec{G}_1(\vec{F}_1(t)) \tag{56}$$

$$\frac{d\ \vec{F}_2(t)}{dt} = \tau\vec{G}_2(\vec{F}_2(t)) \tag{57}$$

#### Figure S1

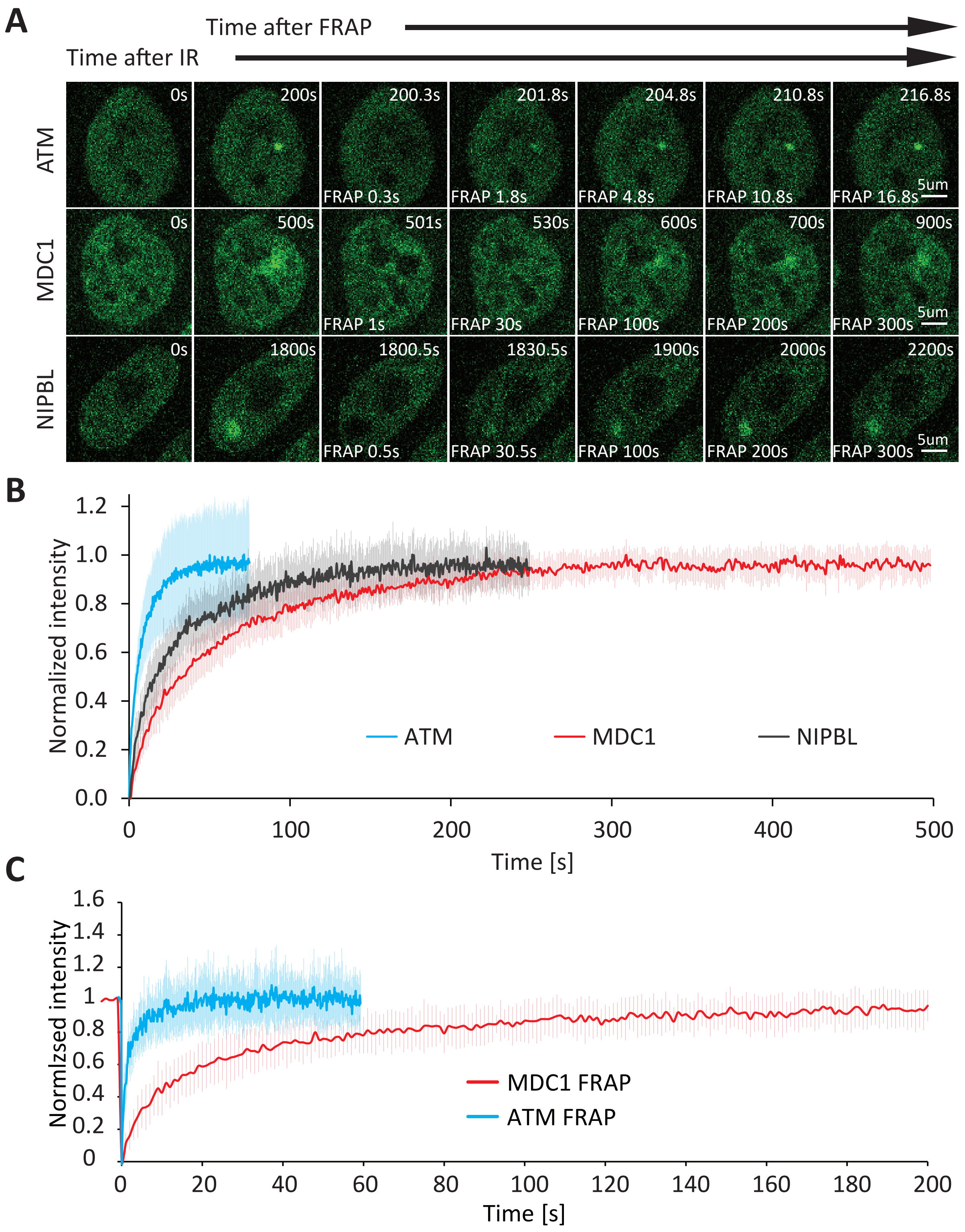


**Figure S1.** FRAP analysis of ATM, MDC1, and mNIPBL foci at micro-irradiation-induced complex DNAlesions.

(A) Representative time-lapse microscopy images of ATM, MDC1, and mNIPBL at complex DNA lesions.

(B) FRAP of micro-irradiation-induced ATM, MDC, and mNIPBL. 0 s corresponds to the first frame after photobleaching. Intensity is normalized to the maximum value after photobleaching.

(C) FRAP of nuclear MDC1 and ATM protein in non-irradiated cells.

#### Figure S2

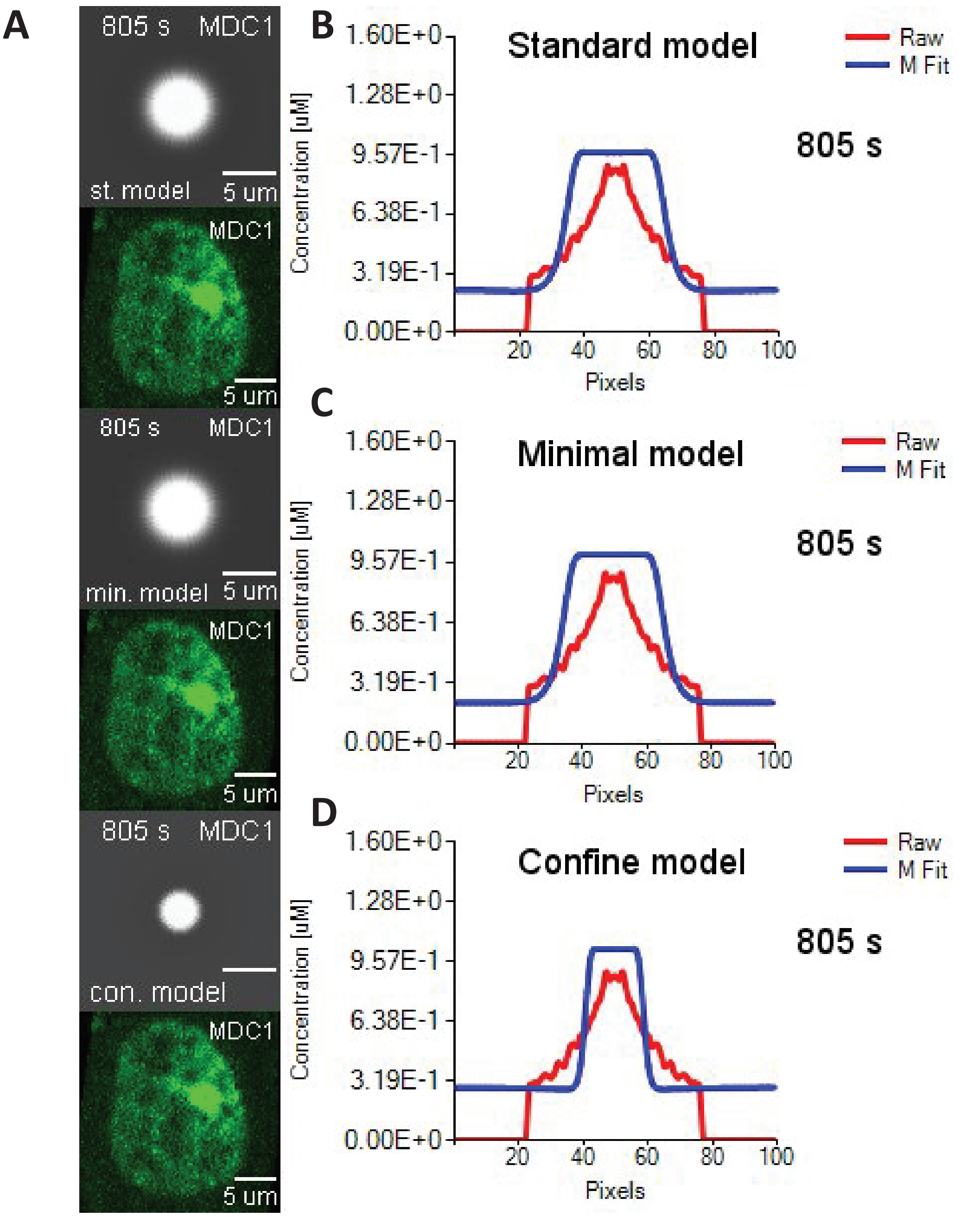


Figure S2. Comparison between the experimentally measured and predicted spatial distribution of MDC1 concentration across the sites of complex DNA damage at 805 s after micro-irradiation. All time points can be observed in Movie S4.

(A) Microscopy images and simulation of MDC1 recruitment based on each model's best fit parameters. Grey images represent the images of MDC1 recruitment generated according to the model. Microscopy images are colored in green.

(B) Comparison between the experimentally measured and predicted (Standard model) spatial distribution of MDC1 concentration across the sites of complex DNA damage at 120 s after micro-irradiation.

(C) Comparison between the experimentally measured and predicted (Minimal model) spatial distribution of MDC1 concentration across the sites of complex DNA damage at 120 s after micro-irradiation.

(D) Comparison between the experimentally measured and predicted (Confined model) spatial distribution of MDC1 concentration across the sites of complex DNA damage at 120 s after micro-irradiation.

### Figure S3

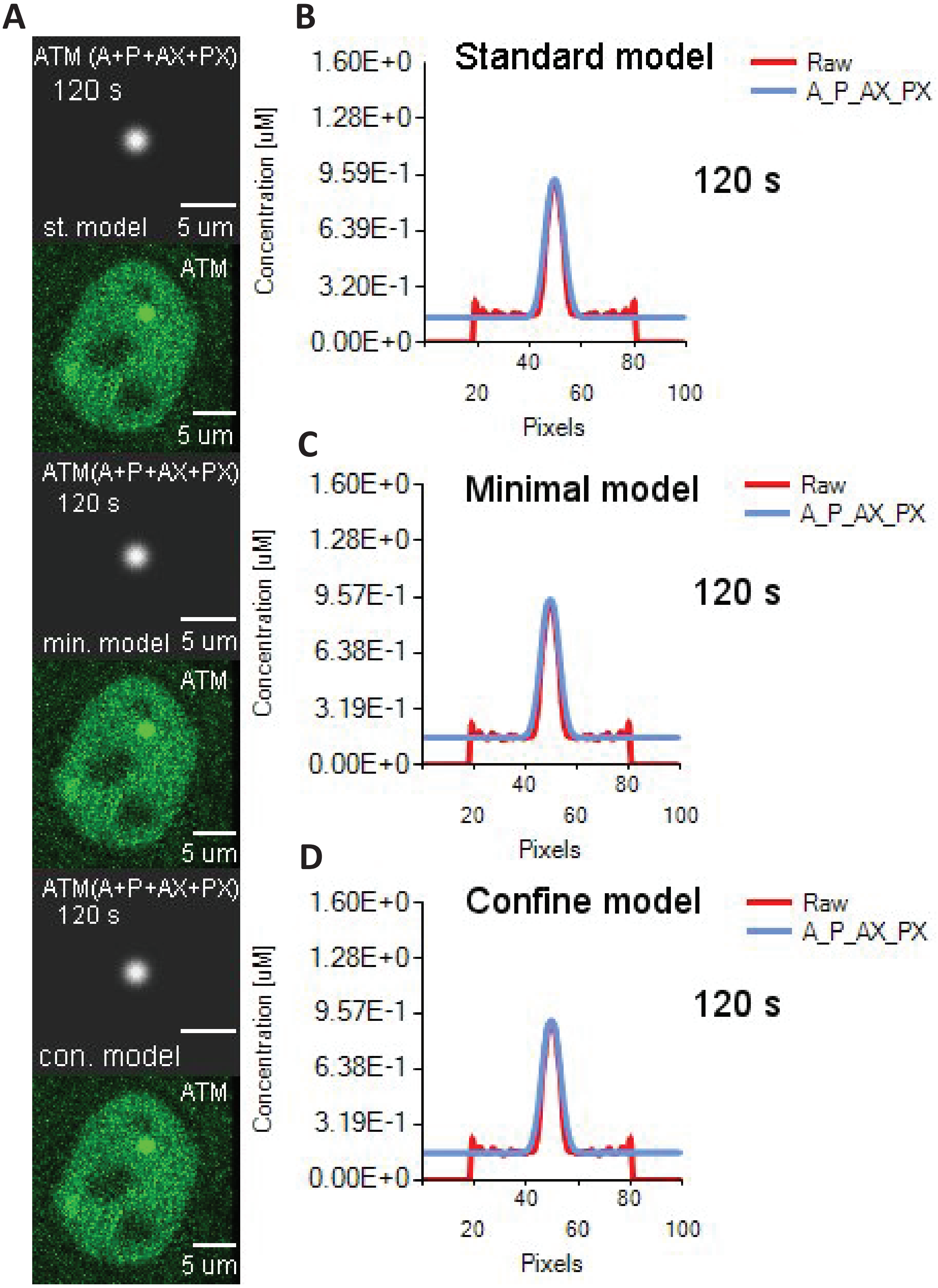
A
ATM (A+P+AX+PX)
120 s
st. model
5 um
ATM
5 um
ATM(A+P+AX+PX)
120 s
min. model
5 um
ATM
5 um
ATM(A+P+AX+PX)
120 s
con. model
ATM
5 um
B
Standard model
Raw
A_P_AX_PX
120 s
Concentration [uM]
1.60E+0
1.28E+0
9.59E-1
6.39E-1
3.20E-1
0.00E+0
20
40
60
80
100
Pixels
C
Minimal model
Raw
A_P_AX_PX
120 s
Concentration [uM]
1.60E+0
1.28E+0
9.57E-1
6.38E-1
3.19E-1
0.00E+0
20
40
60
80
100
Pixels
D
Confine model
Raw
A_P_AX_PX
120 s
Concentration [uM]
1.60E+0
1.28E+0
9.57E-1
6.38E-1
3.19E-1
0.00E+0
20
40
60
80
100
Pixels

**Figure S3.** Comparison between the experimentally measured and predicted spatial distribution of ATM concentration across the sites of complex DNA damage at 120 s after micro-irradiation. All time points can be observed in **Movie S5**.

(A) Microscopy images and simulation of ATM recruitment based on each model's best fit parameters. Grey images represent the images of ATM recruitment generated according to the model. "A" refers to freely diffusing inactive ATM, "AX" refers to inactive ATM bound to the damage site, "P" refers to freely diffusing active ATM (αATM), and "PX" refers to active ATM bound to the damage site. Microscopy images are colored in green.

(B) Comparison between the experimentally measured and predicted (Standard model) spatial distribution of ATM concentration across the sites of complex DNA damage at 120 s after micro-irradiation.

(C) Comparison between the experimentally measured and predicted (Minimal model) spatial distribution of ATM concentration across the sites of complex DNA damage at 120 s after micro-irradiation.

(D) Comparison between the experimentally measured and predicted (Confined model) spatial distribution of ATM concentration across the sites of complex DNA damage at 120 s after micro-irradiation.

#### Figure S4

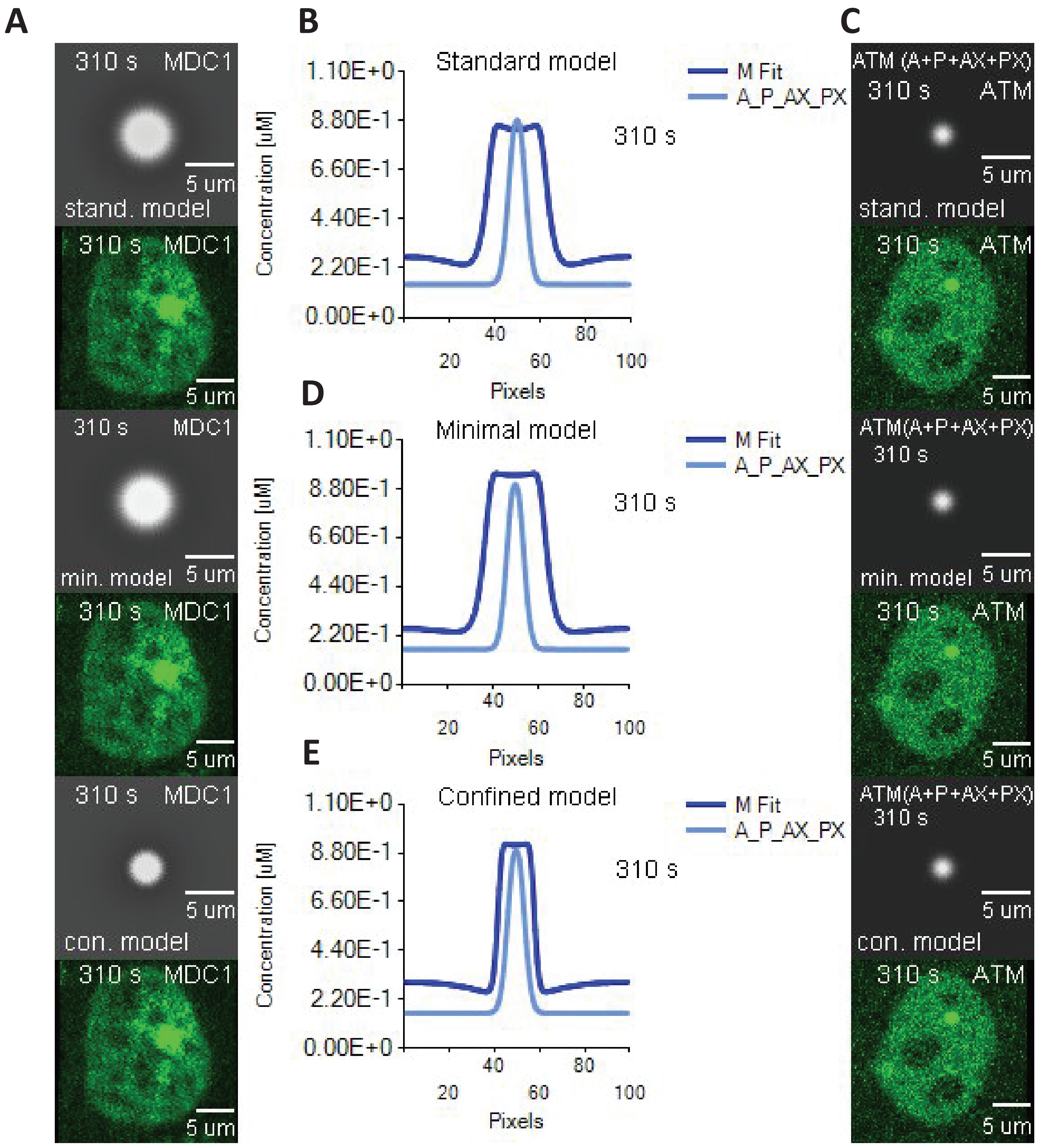
A
B
C
310 s MDC1
5 um
stand. model
310 s MDC1
5 um
310 s MDC1
min. model 5 um
310 s MDC1
5 um
310 s MDC1
5 um
con. model
310 s MDC1
5 um
Standard model
M Fit
A_P_AX_PX
310 s
Concentration [uM]
1.10E+0
8.80E-1
6.60E-1
4.40E-1
2.20E-1
0.00E+0
20 40 60 80 100
Pixels
D
Minimal model
M Fit
A_P_AX_PX
310 s
E
Confined model
M Fit
A_P_AX_PX
310 s
ATM (A+P+AX+PX)
310 s ATM
5 um
stand. model
310 s ATM
5 um
ATM(A+P+AX+PX)
310 s
min. model 5 um
310 s ATM
5 um
ATM(A+P+AX+PX)
310 s
5 um
con. model
310 s ATM
5 um

**Figure S4.** Comparison between the predicted spatial distribution of ATM and MDC1 concentrations across sites of complex DNA damage at 310 s after micro-irradiation. All time points can be observed in **Movie S6.**

(A) Microscopy images and simulation of MDC1 recruitment based on each model's best fit parameters. Grey images represent the images of ATM recruitment generated according to the three models.  Microscopy images are colored in green.

(B) Comparison between predicted spatial distribution of ATM and MDC1 concentration across the sites of complex DNA damage at 310 s after micro-irradiation based on the Standard model.

(C) Microscopy images and simulation of ATM recruitment based on each model's best fit parameters. Grey images represent the images of ATM recruitment generated according to the model.  "A" refers to freely diffusing inactive ATM, "AX" refers to inactive ATM bound to the damage site, "P" refers to freely diffusing active ATM (αATM), and "PX" refers to active ATM bound to the damage site. Microscopy images are colored in green.

(D) Comparison between predicted spatial distribution of ATM and MDC1 concentration across the sites of complex DNA damage at 310 s after micro-irradiation based on the Minimal model.

(E) Comparison between predicted spatial distribution of ATM and MDC1 concentration across the sites of complex DNA damage at 310 s after micro-irradiation based on the Confined model.

#### Figure S5

A

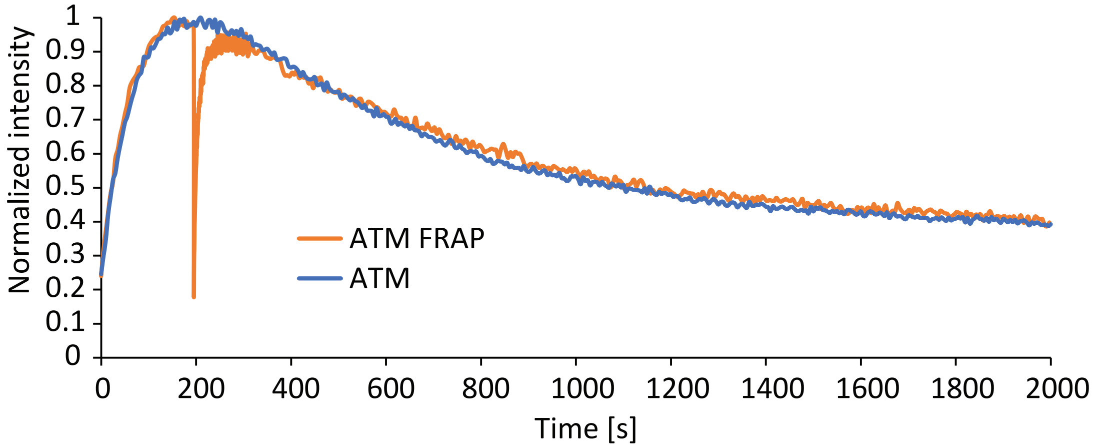


B

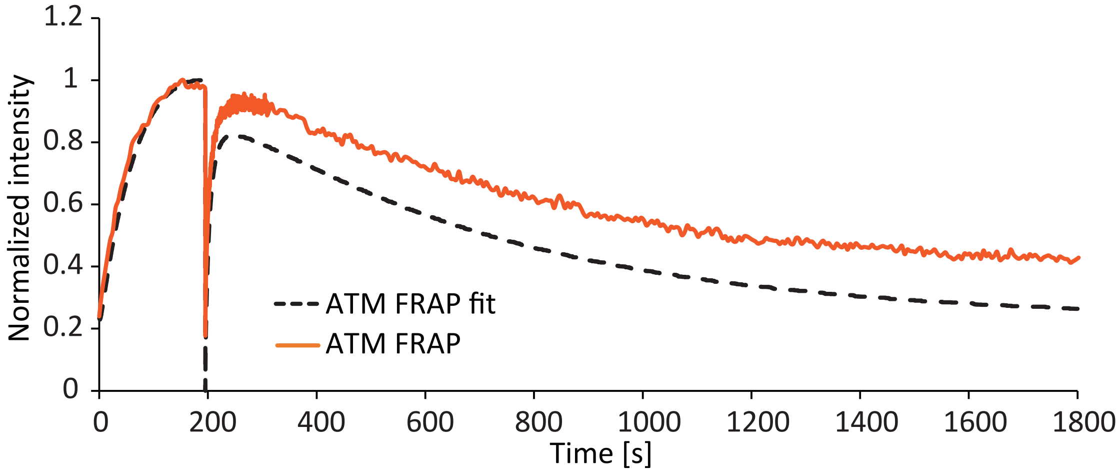


**Figure S5.** Modelling of protein dynamics at micro-irradiation-induced complex lesions after FRAP.

(A) Comparison of ATM dynamics at complex lesions with and without FRAP.

(B) Comparison of measured ATM dynamics at complex lesions after FRAP with those predicted based on the standard model.

### Figure S6

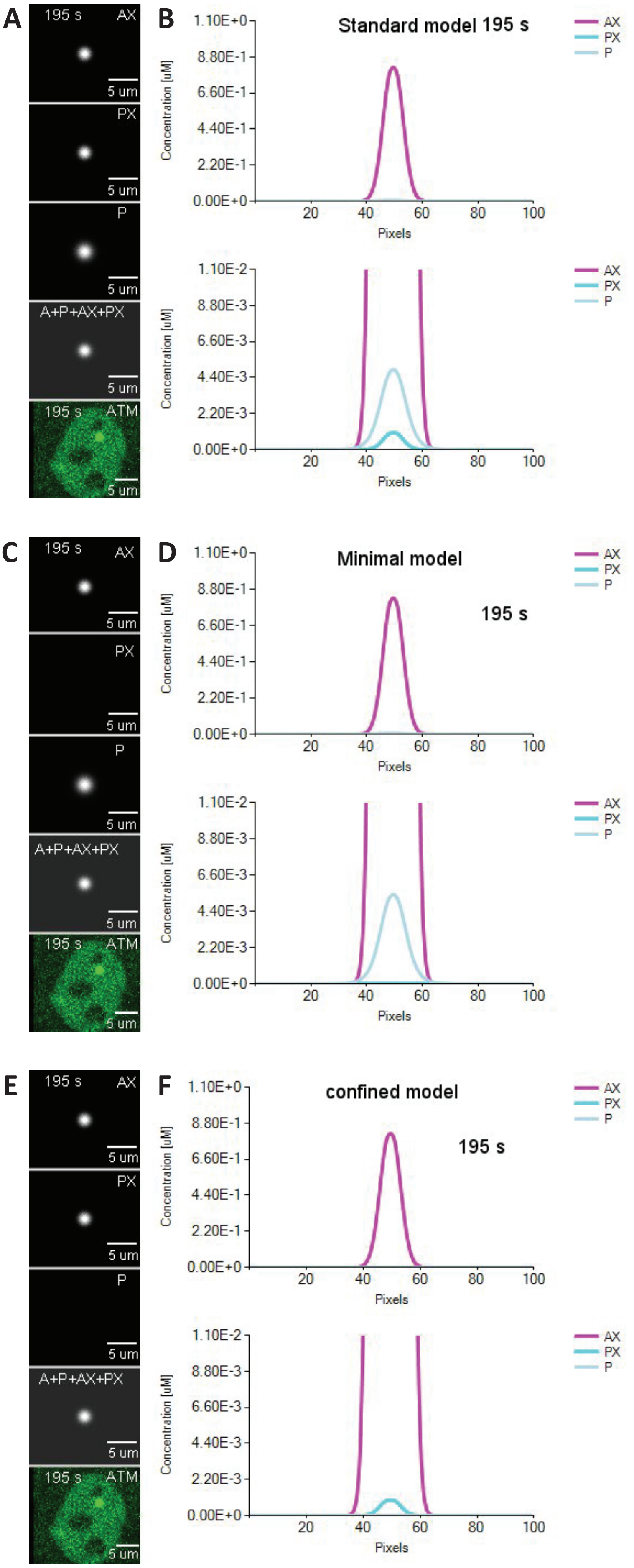
A
195 s
AX
PX
P
A+P+AX+PX
ATM
5 um
B
Standard model 195 s
C
D
Minimal model
195 s
E
F
confined model
195 s
Concentration [uM]
Pixels
1.10E+0
8.80E-1
6.60E-1
4.40E-1
2.20E-1
0.00E+0
1.10E-2
8.80E-3
6.60E-3
4.40E-3
2.20E-3
20
40
60
80
100

**Figure S6.** Comparison between the predicted spatial distribution of active/inactive and bound/freely diffusing ATM populations across the sites of complex DNA damage at 195 s after micro-irradiation. All time points can be observed in **Movie S7**.

(A) Microscopy images and simulation of ATM recruitment based on the Standard model's best fit parameters. Grey images represent the images of ATM recruitment generated according to the model. "A" refers to freely diffusing inactive ATM, "AX" refers to inactive ATM bound to the damage site, "P" refers to freely diffusing active ATM, and "PX" refers to active ATM bound to the damage site. Microscopy images are colored in green.

(B) Comparison between the predicted (Standard model) spatial distribution of AX, PX, and P at the DNA damage site at 195 s after micro-irradiation. The second graph in B is shown on a blown-up scale of the first graph.

(C) Microscopy images and simulation of ATM recruitment based on the Minimal model's best fit parameters. Grey images represent the images of ATM recruitment generated according to the model. "A" refers to freely diffusing inactive ATM, "AX" refers to inactive ATM bound to the damage site, "P" refers to freely diffusing active ATM (αATM), and "PX" refers to active ATM bound to the damage site. Microscopy images are colored in green.

(D) Comparison between the predicted (Minimal model) spatial distribution of AX, PX, and P at the DNA damage site at 195 s after micro-irradiation. The second graph in D is shown on a blown-up scale of the first graph.

(E) Microscopy images and simulation of ATM recruitment based on the Confined model's best fit parameters. Grey images represent the images of ATM recruitment generated according to the model. "A" refers to freely diffusing inactive ATM, "AX" refers to inactive ATM bound to the damage site, "P" refers to freely diffusing active ATM, and "PX" refers to active ATM bound to the damage site. Microscopy images are colored in green.

(F) Comparison between the predicted (Confined model) spatial distribution of AX, PX, and P at the DNA damage site at 195 s after micro-irradiation. The second graph in F is shown on a blown-up scale of the first graph.

**Table S1**

| | Obtained from model | | | Measured |
|---|---|---|---|---|
| | Standard | Minimal | Confined | |
| αATM diffusion coefficient [$\mu m^2 s^{-1}$] | 0.05161 | 0.051613 | | |
| ATM diffusion coefficient [$\mu m^2 s^{-1}$] | 0.663615 | 0.663615 | 0.663615 | 0.6636 ± 0.335 |
| MDC1 effective diffusion coefficient [$\mu m^2 s^{-1}$] | 0.05013 | 0.05133 | 0.03314 | 0.0551 ± 0.0351 |
| $k_0$ [$s^{-1}$] | 0.01368 | 0.012683 | 0.011168 | |
| $k_{on}$ [$M^{-1} \cdot s^{-1}$] | 4.898486 | 4.898486 | 13.00139 | |
| $k_{off}$ [$s^{-1}$] | 0.230875 | 0.230875 | 0.270209 | |
| $k_3$ [$M.s^{-1}$] | 0.002541 | 0.002441 | 0.003062 | |
| $k_d$ [$s^{-1}$] | 0.201373 | 0.191308 | 0.605091 | |
| $k_4$ [$s^{-1}$] | 1.8 | | 2 | |
| $k_{fh}$ [$M^{-1} \cdot s^{-1}$] | 25.02632 | 25.02632 | 60.14428 | |
| $k_{rh}$ [$s^{-1}$] | 0 | 0 | 0 | |
| $k_{on\ MDC1}$ [$M^{-1} \cdot s^{-1}$] | 0.488357 | 0.488357 | 0.488357 | |
| $k_{off\ MDC1}$ [$s^{-1}$] | 0.109351 | 0.109351 | 0.109351 | |
| X [μM] | 1.452281 | 1.492281 | 1.342281 | |
| ATM concentration [μM] | 0.155023 | 0.155023 | 0.155023 | 0.155023 |
| H2AX concentration [μM] | 1.59275 | 1.59275 | 1.39275 | |
| MDC1 concentration [μM] | 0.302879 | 0.302879 | 0.302879 | 0.302879 |

**Table S1** Rate constants, diffusion coefficients and concentrations of reactants of the reactions 1-19 that was measured during the experiments or obtained by fitting the mathematical models to experimental data.
